# An Aphid Protein Interferes with Plant endomembrane Trafficking Machinery to Promote Aphid Survival and Plant Growth

**DOI:** 10.64898/2026.10.02.756256

**Authors:** Leila Feiz, Navid Movahed, Honglin Feng, Venkatesh Thirumalaikumar, Sergey Ivanov, Devender Arora, Jander Georg

**Affiliations:** BOYCE Thompson Institute, Ithaca, NY 14853, USA; Department of Entomology, Louisiana State University; Purdue Proteomics Facility, Bindley Biosceince Center, Purdue University; Center for Plant Biology, Purdue University; Department of Botany and Plant Pathology Oregon State University.

## Abstract

Aphids deplete their host plants of amino acids and other essential nutrients by feeding on phloem sap. Autophagy, a conserved process across eukaryotes, maintains cellular homeostasis during starvation and environmental stress by recycling damaged or excess cellular components. Here, we identified and characterized a previously uninvestigated aphid salivary protein, termed Aphid Autophagy-Related Factor 1 (AAF1), that is secreted into plant cells and manipulates plant physiology in a manner that benefits the aphid. We show that AAF1 interacts with four known or putative host autophagy-related proteins: RABD1 and RABD2a (plant homologs of Ypt1/RAB1), ATG8, a ubiquitin-like protein required for autophagosome formation, and PI3PT, a previously uncharacterized ER-localized putative lipid-binding or lipid-transfer protein annotated as a phosphoinositide 3-phosphate transfer protein. We further demonstrate that the RAB-binding domain of AAF1 and its interaction with RABD2a may be specifically required for AAF1 function. In addition, we identified unique, highly mobile compartments containing interacting AAF1 and RABD2a that are associated with the endoplasmic reticulum (ER) and dynamically traffic with autophagosomes. We propose that these compartments may be derived from endosomes or endosome–ER contact sites (EERCSs) and serve as platforms for autophagosome trafficking and/or biogenesis. We further propose that AAF1, RABD2a, and PI3PT function together to regulate the formation or function of these compartments, thereby supporting autophagosome formation and trafficking. Finally, we show that stable expression of AAF1 in plants enhances growth and biomass in an autophagy-dependent manner.

## Introduction

The interaction between plants and phytopathogens, in which both partners manipulate the physiology of the other, is a central theme in plant biology. Like microbial pathogens, aphids secrete salivary proteins into plant cells to alter host physiology and suppress defense responses, although the functions of most of these proteins remain unknown ^1–7^. Here, we characterize one such previously uninvestigated aphid salivary protein that interacts with the plant vesicle trafficking machinery and manipulates autophagy.

Macroautophagy (hereafter autophagy) is a conserved eukaryotic pathway that delivers cytoplasmic components to the lytic compartment for degradation through double-membrane autophagosomes ^8–11^. In plants, autophagy is activated by nutrient limitation and environmental stresses and plays essential roles in development, nutrient recycling, immunity, and stress adaptation ^12–27^.

Autophagosome biogenesis is mediated by more than 40 conserved autophagy-related (ATG) proteins that assemble at the pre-autophagosomal structure, or phagophore assembly site (PAS), to coordinate phagophore initiation and expansion ^9,28,29^. Under starvation conditions, activation of the ATG1 complex recruits the phosphatidylinositol 3-kinase complex (PI3KC3-C1), leading to the production of phosphatidylinositol 3-phosphate (PI3P) on ATG9-positive membranes derived from the ER, Golgi, plasma membrane, and endosomal compartments. PI3P-binding proteins subsequently recruit downstream factors, including the ATG12–ATG5–ATG16 and ATG2 complexes, to promote ATG8 lipidation and membrane expansion ^29–33^. During selective autophagy, cargo receptors interact with the ATG1 complex, ATG11, and lipidated ATG8 to couple cargo sequestration with phagophore growth ^34–41^.

Membranes supporting autophagosome expansion are supplied by multiple endomembrane compartments, including ER-derived Coat Protein Complex II (COPII) vesicles and Golgi- or endosome-derived ATG9 vesicles ^42–44^. The recruitment of COPII vesicles to the PAS is coordinated by the Rab GTPase Ypt1 and its mammalian and plant homologs, RAB1 and RABD2a, respectively ^45–50^.

Many of the factors involved in autophagosome biogenesis that are conserved across eukaryotes have been characterized in plants ^11,12,45,51–55^. Diverse phytopathogens have also evolved effectors that manipulate host autophagy to promote infection, yet relatively few of these effectors have been functionally characterized ^56–61^. One notable example is the *Phytophthora infestans* effector PEXRD54, which redirects selective autophagy by binding ATG8CL and recruiting lipid droplets through RAB8a to support autophagosome formation ^57–59^.

Here, we identify and characterize a previously uninvestigated aphid salivary protein, Aphid Autophagy-Related Factor 1 (AAF1), which interacts with multiple host proteins, including RABD2a, a protein associated with Golgi stacks and trans-Golgi– endosomal compartments ^62–64^ that is involved in vesicle trafficking and autophagy ^45^. Together, AAF1 and RABD2a form a complex on a highly dynamic, ER-associated compartment that likely generates and/or traffics autophagosomes. We also show that constitutive expression of AAF1 enhances Arabidopsis growth and biomass through an autophagy-dependent mechanism.

## Results and Discussion

### AAF1, an Aphid Salivary Protein, Functions in Plant–Aphid Interactions

In a broad search for aphid-secreted proteins, we conducted a proteomic analysis of artificial diet that had been fed upon by *Myzus persicae* (green peach aphids). From thousands of salivary proteins detected in this assay, 120 were predicted to contain a secretion signal peptide. Genes encoding 44 small-sized salivary proteins (10–25 kDa, excluding their signal peptides) (Fig. S1a) were cloned into high-expression pEAQ vectors ^65^, with an EGFP tag (referred to here as GFP for simplicity) fused to either the N-or C-terminus under the control of the 35S promoter. These constructs were introduced into *Agrobacterium tumefaciens* for transient expression in *Nicotiana benthamiana* leaf cells (Fig. S1b) to assess their effects on leaf symptoms, aphid reproduction, and subcellular localization. Prolonged overexpression of one of these unknown proteins, AAF1, caused cell death like symptoms on the leaf (GFP-AAF1 in Fig. 1a). The cell death effect caused by AAF1 in leaves was alleviated when the protein was expressed at lower levels using an alternative vector (AAF1-HA(3x) and HA-AAF1 in Figs. 1a, S2a), as determined by immunoblot analysis with an antibody raised against the protein (Fig. S2b). The AAF1 protein is encoded by *M. persicae* gene *MYZPE13164_G006* (https://bipaa.genouest.org/is/aphidbase/; GenBank *LOC111028600*), which is conserved in other genome-sequenced aphid species (Fig. S3), but is not found in other organisms, including species in the closely related Adelgidae and Phylloxeridae. Aphid genomes contain several hundred genes encoding proteins not found anywhere else in the kingdom of life. The emergence of new protein-coding genes, likely arising de novo from non-coding DNA sequences in aphids, is not unique to this taxon. Recent advances in genome and transcriptome sequencing, along with ribosome profiling, have uncovered instances of de novo gene birth across a wide range of organisms, from yeast to rice ^66,67^. A newly born gene can become fixed in a population if it is subject to positive selection because the protein it encodes provides a fitness advantage to the host organism or improves its interaction with the environment ^68^.

**Fig. 1.**
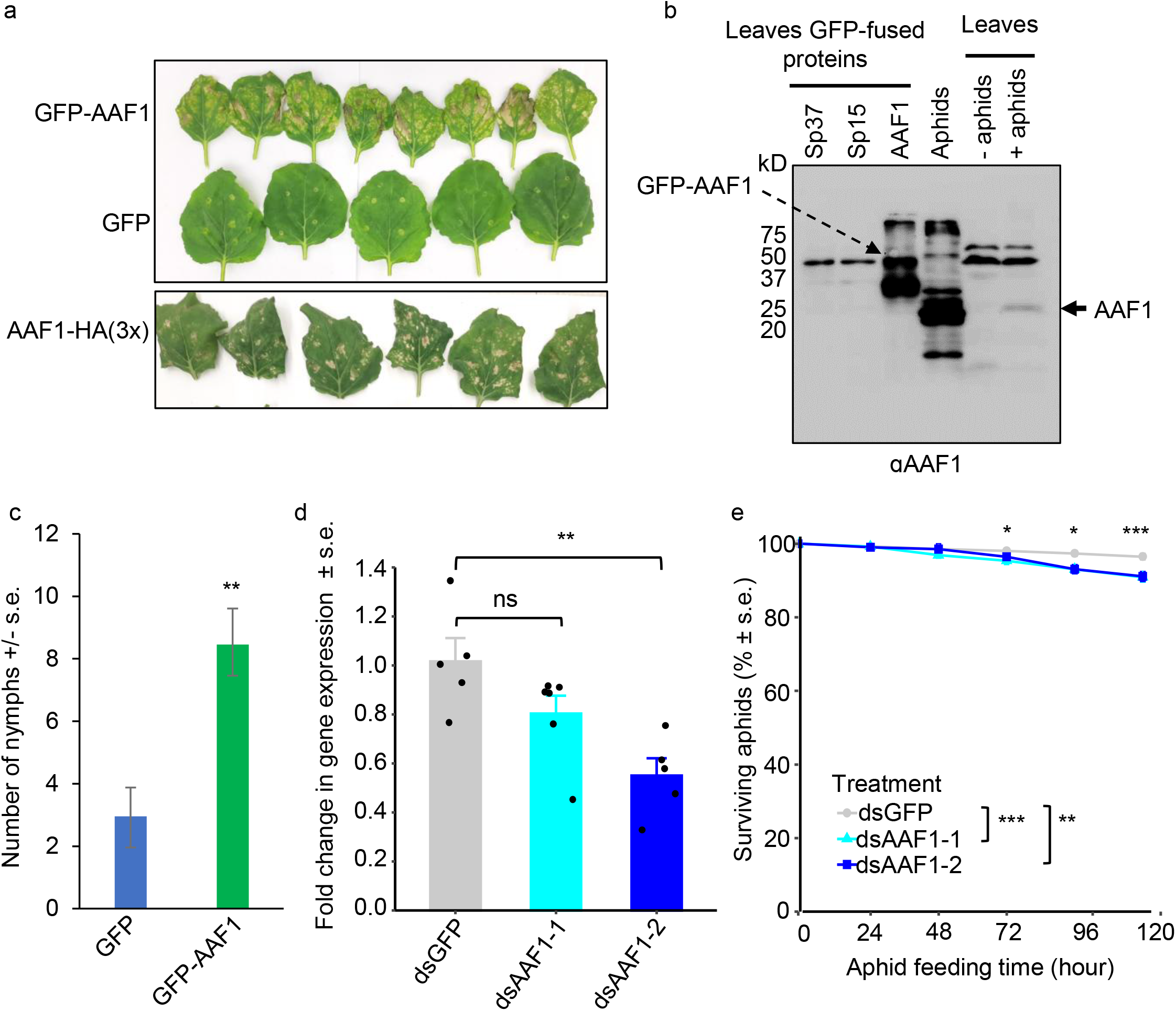
AAF1 functions in plant–aphid interactions. (**a**) Transient expression of AAF1 induces cell death-like symptoms in *N. benthamiana*. GFP-AAF1, GFP, and AAF1-HA(3×) were transiently expressed and imaged 10 days after infiltration. (**b**) Aphids secrete AAF1 into plant leaves during feeding. To assess AAF1 presence, immunoblot analysis was conducted on total protein extracts from *N. benthamiana* leaves transiently expressing GFP fusions of AAF1, Sp15, and Sp37 (the latter two serving as negative controls), from *M. persicae* aphid bodies, from a control *N. tabacum* leaf without aphids, and from a *N. tabacum* leaf collected one day after aphid feeding, following aphid removal and leaf washing. Protein blots were probed with antibodies specific to AAF1. A few nonspecific bands comigrated with or migrated above GFP-AAF1 and were also present in samples lacking AAF1. (**c**) Transient expression of GFP-AAF1 in *asat2 N. benthamiana* increased aphid reproduction compared with GFP. Data are mean ± s.e. (N = 24). \*\**P* < 0.01 (t test). Raw data are provided in Data Set S1. (**d**) Plant-mediated virus-induced gene silencing (VIGS) was used to knock down *AAF1* in aphids using two independent constructs (ds-*AAF1-1* and ds-*AAF1-2*). (**e**) *AAF1* knockdown reduced aphid survivorship relative to the ds-*GFP* control. Data were collected from 5–7 plants with two cages per plant. Statistics were analyzed by ANOVA with block effects followed by Dunnett’s test. ns, not significant; \*\**P* < 0.01; \*\*\**P* < 0.001.

Since AAF1 was initially identified only in aphid saliva mixed with an artificial diet, we sought to confirm its secretion into plant tissue. Immunoblot analysis of total leaf protein extracts showed that the antibody recognized both native AAF1 in aphid bodies and GFP-tagged AAF1 transiently expressed in *N. benthamiana* (Fig. 1b). As negative controls, protein extracts were prepared from leaves transiently expressing two unrelated GFP-tagged aphid proteins, SP15 and SP37 (Fig. S1b). Additionally, the antibody detected a band corresponding to AAF1 in *Nicotiana tabacum* (tobacco) leaves that had been exposed to aphids for one day, but not in control leaves without aphid exposure (Fig. 1b). In the immunoblot, several bands corresponding to non-specific antibody binding comigrated with or migrated above GFP–AAF1 and were also present in lanes containing samples lacking GFP–AAF1. The presence of aphid-secreted AAF1 in leaves was further confirmed by immunoprecipitation of native AAF1 from aphid-fed leaves followed by LC–MS analysis.

To determine whether AAF1 enhances aphid performance or plant resistance, we conducted a fecundity bioassay using *N. benthamiana* leaves transiently expressing GFP-AAF1 or GFP (negative control). Aphids reproduced significantly more on AAF1-expressing leaves, indicating that AAF1 benefits aphids (Fig. 1c). Because hypersensitive response (HR)-associated programmed cell death and reactive oxygen species (ROS) accumulation typically enhance plant defense against bacterial pathogens, we examined whether AAF1 induces hydrogen peroxide (H₂O₂), a major ROS that accumulates during aphid infestation ^69^. We confirmed that AAF1 induces H₂O₂ accumulation (Fig. S4a). Coexpression of Catalase1, a peroxisomal enzyme that scavenges H₂O₂, reduced both H₂O₂ levels and cell death symptoms (Figs. S4a and b). In addition, Catalase1 overexpression reduced aphid fecundity and survivorship (Figs. S4c and d). Together, these findings indicate that H₂O₂ accumulation and associated cell death do not enhance resistance to aphids and may instead promote aphid performance. Unlike bacterial pathogens, whose spread can be restricted by localized cell death, aphids can move freely across plant tissues, making such responses ineffective as a defense strategy. To further support the conclusion that aphids secrete AAF1 into plant tissues to promote their own fitness, we used plant-mediated virus-induced gene silencing (VIGS) ^70^ to knock down *AAF1* expression and compared the survivorship of adult aphids with reduced *AAF1* transcript levels on their *N. benthamiana* host plants. Consistent with the fecundity bioassay, two independent *AAF1* RNAi knockdown aphid lines exhibited significantly reduced survivorship compared with the *GFP* RNAi control (Figs. 1d, e, and S5).

AAF1 was recently identified in the watery saliva of the grain aphid *Sitobion miscanthi*, which was collected from *Triticum aestivum* (bread wheat), and its transcript (*Sm13498*) was detected in the salivary glands of the aphid ^71–73^. Notably, the transcript level was highest in apterous *S. miscanthi* adults after feeding on wheat leaves for 12 hours ^73^. Together, these findings support our conclusion that AAF1 plays a conserved and essential role in plant-aphid interactions.

### AAF1 Interacts with RABD1 and RABD2a via a Rab-Binding Domain

To further investigate the role of AAF1, we identified interacting plant proteins by performing GFP trap-mediated co-immunoprecipitation (Co-IP) of GFP–AAF1 from *N. benthamiana* leaves transiently expressing the protein, followed by LC–MS analysis of co-purifying host proteins. To maximize protein identification, LC–MS data were searched against a combined *N. benthamiana* and *N. tabacum* protein database because the *N. tabacum* proteome is more complete and the two species share high genomic and proteomic similarity. Across two independent experiments, each comprising multiple biological replicates, AAF1 consistently interacted with three plant proteins: the closely related Rab GTPases RABD1 and RABD2a, and an uncharacterized lipid-binding protein discussed in a later section. RABD1 and RABD2a are conserved plant homologs ^45,62^ of mammalian RAB1 and yeast Ypt1, Rab GTPases with well-established roles in ER-to-Golgi trafficking, endosome dynamics, and autophagosome biogenesis ^42,47,48,50,74–76^. In *N. tabacum*, RABD1 (XP_016461491) and RABD2a (XP_016478812/XP_016495270) are encoded by *LOC107784791* and *LOC107814370*, respectively. The corresponding *N. benthamiana* homologs are encoded by *Niben044Scf00004703:32370* and *Niben101Scf01188g09003.1*, whereas the Arabidopsis homologs are encoded by *AT3G11730* (RABD1) and *AT1G02130* (RABD2a).

RAB proteins, including RAB1, play essential roles in endosomal trafficking and maturation ^76–78^. In yeast and mammalian cells, Ypt1 and RAB1, respectively, bridge ER-derived COPII vesicles and the associated TRAPPIII complex with the ATG1 kinase complex, a major autophagy-initiation machinery, at the PAS, thereby promoting phagophore formation ^42,47,48,50,74,75,79^. Similarly, in plants, RABD proteins, including RABD1 and RABD2a, localize to punctate structures identified as the trans-Golgi network/early endosome (TGN/EE), where they likely regulate endosomal trafficking ^62–64^. Like its yeast and mammalian counterparts, Arabidopsis RABD2a has been shown to be involved in autophagosome formation ^45^. Overall, studies across yeast, mammals, and plants suggest that the role of RAB1 orthologs in autophagy may be derived from or integrated with their broader functions in ER–endosome trafficking. More recently, recycling endosomes and ER–endosome contacts in mammalian cells have been proposed to serve as platforms for autophagosome biogenesis ^80–83^, further linking RAB1-mediated vesicle and endosomal trafficking to the early stages of autophagosome formation.

We validated the interaction between AAF1 and *Nt-*RABD1 or *Nt*-RABD2a (hereafter referred to as RABD1 and RABD2a) using reverse Co-IP. GFP-tagged RABD1 or RABD2a, used as bait, successfully pulled down N- or C-terminally tagged AAF1 variants, including Flag-AAF1 (Fig. 2a), AAF1-HA(3x), and AAF1-Flag (Fig. S6a and b). In these experiments, GFP-tagged ATG8f, an autophagosome-associated ATG8 isoform, was used as a bait control and showed no detectable interaction with AAF1, similar to the negative control using free GFP. We next examined the subcellular colocalization of RABD1 and RABD2a with AAF1 in *N. benthamiana* leaf cells. Z-stack and single-plane imaging revealed that RFP–RABD2a and GFP–AAF1 colocalize in cytosolic puncta consistent with the trans-Golgi network/early endosome (TGN/EE) structures observed in previous studies ^62–64^ (inset in Fig. 2b, filled arrowhead), some of which are associated with the perinuclear ER (inset in Fig. 2b, open triangle). Similar puncta were observed upon coexpression of RFP–RABD1 and GFP–AAF1 (Fig. S6c). Prolonged overexpression of GFP–RABD1 or GFP–RABD2a in *N. benthamiana* also induced cell death symptoms resembling those triggered by GFP–AAF1 overexpression in leaves (Fig. 2c).

**Fig. 2.**
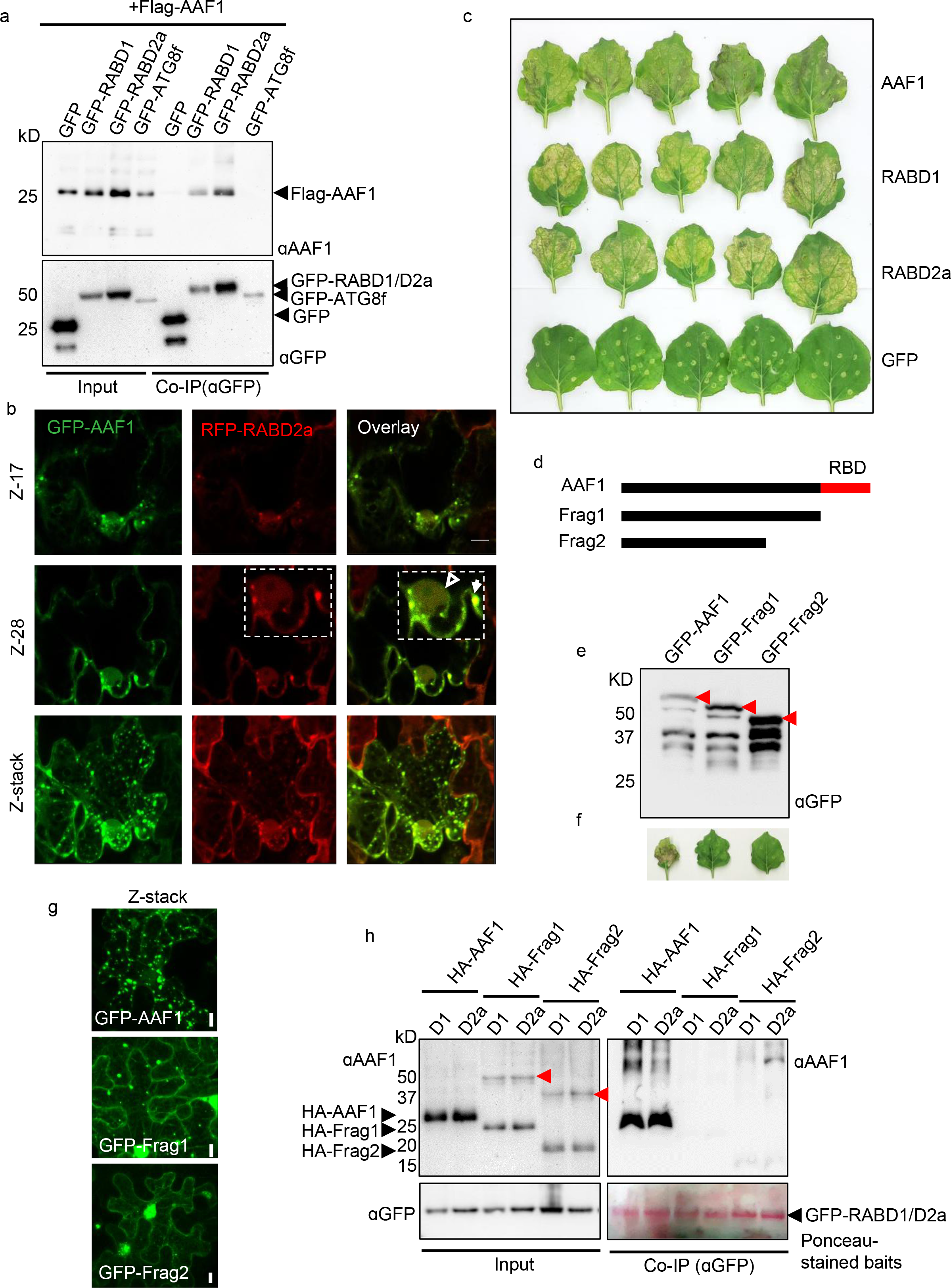
RABD1 and RABD2a associate with AAF1, and AAF1 function depends on its Rab-binding domain. (**a**) GFP or GFP-fused RABD1, RABD2a, or ATG8f were used as baits in GFP-Trap Co-IP of Flag-AAF1 coexpressed with each bait in *N. benthamiana*. Co-IP eluates and input samples were immunoblotted with anti-AAF1 and anti-GFP antibodies. (**b**) Confocal images showing colocalization of GFP-AAF1 (green) and RFP-RABD2a (red) in *N. benthamiana* leaves. Two z-stack sections and the full z-stack are shown. Insets are shown at 2× magnification. Filled arrowhead, punctate structure; open triangle, endoplasmic reticulum surrounding the nucleus. Scale bar, 10 µm. (**c**) Overexpression of RABD1 or RABD2a induces cell death-like symptoms similar to those caused by AAF1. Leaves were imaged 10 days after infiltration. (**d**) Schematic of full-length AAF1 (25.7 kDa) and truncated fragments (Frag1, 23.6 kDa; Frag2, 14.8 kDa; Frag3, 9.12 kDa). (**e**) Anti-GFP immunoblot of total proteins from *N. benthamiana* leaves expressing GFP-AAF1 or GFP-tagged AAF1 fragments 2 days after infiltration. Red arrows indicate the expected GFP fusion proteins. (**f**) Leaf symptoms 10 days after infiltration in leaves expressing GFP-AAF1 or its truncated fragments. (**g**) Confocal z-stack images of GFP-AAF1 and its truncated fragments 2 days after infiltration. Scale bar, 10 µm. (**h**) GFP-RABD1 or GFP-RABD2a were used as baits in GFP-Trap Co-IP assays with HA-tagged full-length AAF1 or truncated fragments (Frag1 and Frag2). Co-IP eluates and input samples were immunoblotted with anti-AAF1 and anti-GFP antibodies. The blot corresponding to the GFP-RABD1/RABD2a Co-IP eluates was stained with Ponceau S instead of being probed with anti-GFP antibody. Red arrows indicate SDS-resistant higher-order species of HA-Frag1 and HA-Frag2, possibly ubiquitinated forms or AAF1 intermediates.

To identify AAF1 domains that may interact with RABD1/RABD2a, we used Phyre2 ^84^ and I-TASSER ^85^ structural modeling. Full-length AAF1 modeling with Phyre2 did not identify consistent candidates with sufficient confidence. We therefore divided AAF1 into six segments (Fig. S7a) and modeled each individually. Both platforms identified a C-terminal region (amino acids 220–250) with >50% structural similarity to domains of the Mon1–Ccz1 complex, a guanine nucleotide exchange factor (GEF) for Ypt7/RAB7 in yeast ^86,87^ and a related human RAB GEF complex ^88^ (Fig. S7). Specifically, Phyre2 identified this region using Segment 6, whereas I-TASSER identified it using full-length AAF1. Phyre2 modeling of the remaining segments identified significant structural homology in segments 2–4 (Fig. S7a). Segment 2 showed homology to an uncharacterized protein (76.3% confidence), segment 3 to the tethering (protein-binding) domain of GET5 (Guided Entry of TA) ^89^ (86.9% confidence), and segment 4 to B-cell lymphoma 2 (Bcl-2), a known regulator of apoptosis and autophagy ^90^ (44.3% confidence). Segments 1 and 5 showed no significant homology to known structures. The functional roles of these AAF1 regions remain to be determined.

To assess the functional role of the predicted Rab-binding/activating domain (RBD) in AAF1, we generated sequential truncations lacking the RBD or a larger portion of AAF1 (Frags 1–2), fused these variants to GFP, and transiently expressed them in *N. benthamiana* leaves (Figs. 2d–g). GFP–AAF1 lacking the RBD (Frag1), as well as the shorter variant Frag2, failed to induce cell death symptoms (Fig. 2f). Moreover, the TGN/EE puncta ^62–64^ formed by full-length GFP–AAF1 were markedly reduced in Frag1 and Frag2 (Fig. 2g). Instead, these truncated variants exhibited diffuse localization throughout the cytosol and around the nucleus, presumably associated with the perinuclear ER. Closer examination of GFP–AAF1 lacking the RBD (Frag1), co-expressed with mCherry–HDEL as an ER marker, revealed that, unlike the TGN/EE puncta formed by full-length GFP–AAF1 (Figs. 2b, 2g, and S8a), the truncated protein localized predominantly to punctate structures that appeared to aggregate at the perinuclear ER (Fig. S8b). These findings suggest that the AAF1 Rab-binding domain may be required for its colocalization with RABD on cytosolic TGN/EE puncta.

To further investigate the role of the predicted RBD in mediating the interaction between AAF1 and RABD1 or RABD2a, we co-expressed HA-tagged full-length AAF1 or its truncation variants (Frag1 and Frag2) with GFP–RABD1 or GFP–RABD2a, followed by Co-IP using the RABD proteins as bait. The results showed that only full-length AAF1 interacted with RABD1 and RABD2a (Fig. 2h). Immunoblot analysis of the same Co-IP samples using an anti-ubiquitin antibody revealed abundant ubiquitinated protein species larger than 50 kDa, likely corresponding to GFP–RABD1 and GFP– RABD2a (Fig. S8c). Notably, a fraction of the expressed HA–Frag1 and HA–Frag2 formed SDS-resistant, higher-order protein species in the input samples, which we speculate represent ubiquitinated forms or intermediate adducts of AAF1 (red arrows, Fig. 2h). These species may be unable to undergo proper processing because the truncated proteins lack domains required for interaction with RAB proteins or other binding partners. As mentioned above, ubiquitination and the autophagy machinery are linked through multiple layers of regulation ^91^. The precise nature of the AAF1-containing adducts, as well as the identity of the putative ubiquitinated RABD protein species, remains to be determined.

### AAF1 May Affect Plant Autophagy in a RABD2a-Dependent Manner

Because AAF1 interacts with RABD proteins, shares their subcellular localization, and induces similar leaf symptoms, and because RABD2a functions in vesicle trafficking pathways required for autophagosome formation, we investigated whether AAF1 is associated with or regulates plant autophagy. As an initial step, we examined potential interactions with ATG8, a family of ubiquitin-like proteins that serve as essential structural and functional components of autophagosomes in all eukaryotes. However, ATG8 peptides were not detected in our proteomic analysis of the AAF1 interactome. This was not unexpected because, aside from the ATG12–ATG5 conjugation system, which mediates ATG8 lipidation ^9,29,31^, most autophagy and PAS-associated complexes are not thought to interact directly with ATG8. Instead, ATG8 primarily binds cargo receptors that reside on autophagosomal membranes and link specific cargo to lipidated ATG8 on the expanding autophagosome ^34–41^. AAF1 is unlikely to function as a cargo receptor, as it localizes predominantly to cytosolic puncta 1–5 µm in diameter (Figs. 2, and S8). By contrast, typical cargo receptors accumulate in both cytosolic autophagosomes (0.5–2 µm) and mobile autophagic bodies (<1 µm) within the vacuole during autophagy induction.

To determine whether AAF1 nevertheless associates with ATG8, we tested three *N. tabacum* ATG8 isoforms in pairwise Co-IP assays with AAF1 (Fig. 3a). Among the three isoforms, ATG8d, the *N. tabacum* ortholog of Arabidopsis ATG8c and potato ATG8CL ^57^ exhibited the strongest interaction with AAF1. However, this interaction was considerably weaker than that observed with RABD, which served as a positive control (Fig. 3a). No interaction was detected between AAF1 and ATG8c, and only a faint AAF1 band was observed when GFP–ATG8f was used as bait, consistent with our previous Co-IP experiments, which showed no interaction between AAF1 and ATG8f (Figs. 2a, S6a, and S6b). Cargo specificity in selective autophagy may be determined by distinct ATG8 isoforms ^92^. Because ATG8d is the ortholog of potato ATG8CL—the specific isoform targeted by the *Phytophthora infestans* effector PEXRD54, which substitutes for the plant cargo receptor NBR1 and reprograms host selective autophagy ^57,58,93^—the observed AAF1–ATG8d interaction raises the possibility that AAF1 also functions in selective autophagy. ATG8-binding cargo receptors contain a consensus AIM/LIR motif (W/Y/F–X–X–L/I/V), although variations are common depending on protein context and ATG8 isoform specificity. We identified a putative AIM motif in AAF1 (Fig. S3, red box), which is highly conserved across most aphid species, with limited variation in a few cases. Such degeneration is not unusual if the motif contributes to ATG8 binding, as previously observed for PEXRD54 variants, where AIM degeneration has been linked to regressive effector evolution following host shifts ^94^.

**Fig. 3.**
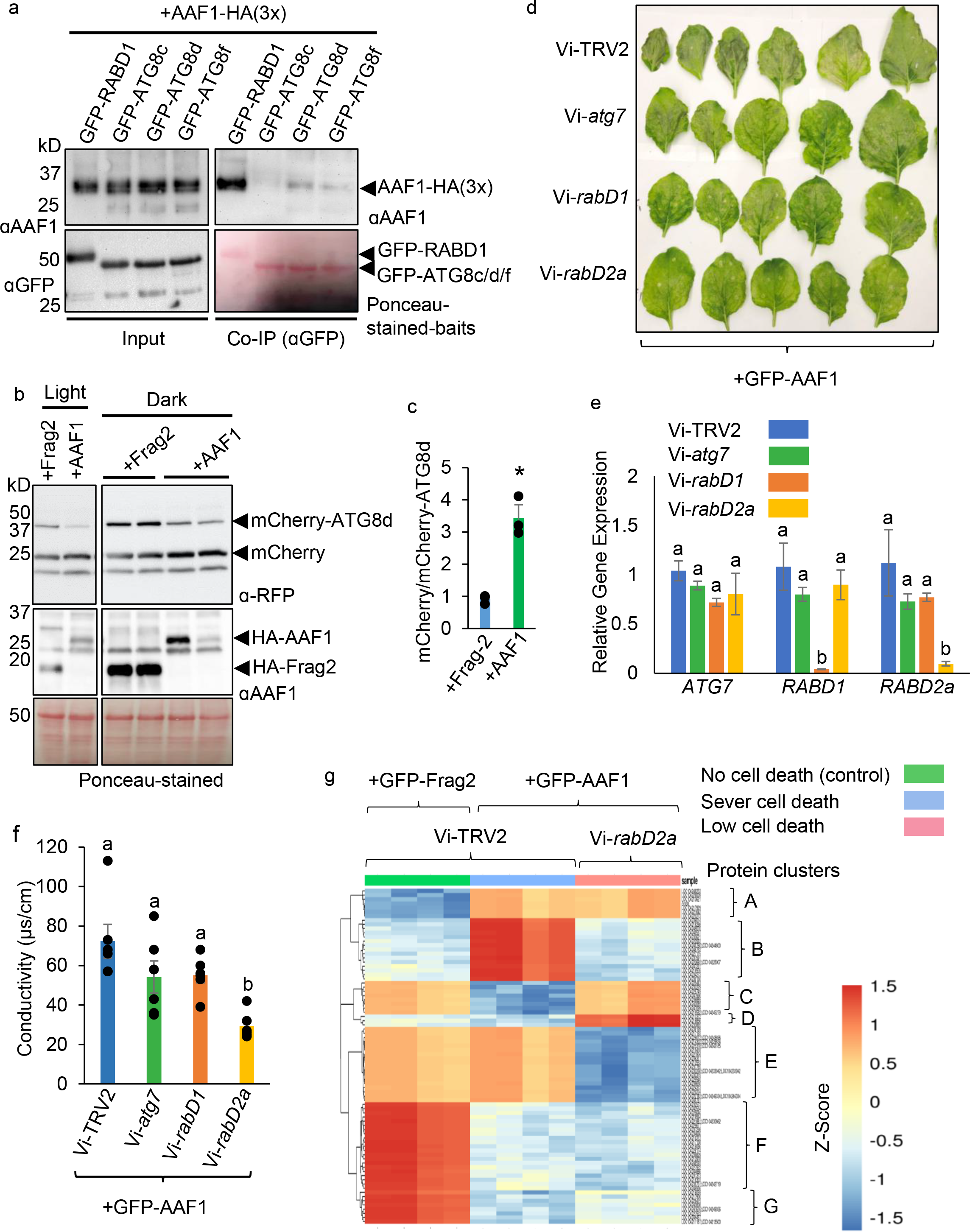
AAF1 is involved in plant autophagy. (**a**) GFP-Trap Co-IP assays were performed using GFP-tagged ATG8c, ATG8d, and ATG8f as baits to pull down HA-AAF1 from *N. benthamiana* leaves coexpressing each GFP-ATG8 with HA-AAF1. GFP-RABD1 served as a positive control. Co-IP eluates and input samples were immunoblotted with anti-AAF1 and anti-GFP antibodies. The blot corresponding to the Co-IP eluates of the GFP-fused baits was stained with Ponceau S rather than being probed with anti-GFP antibody. (**b**) AAF1 increases autophagic flux. Proteins from *N. benthamiana* leaves coexpressing HA-AAF1 or HA-Frag2 with mCherry-ATG8d were immunoblotted with anti-mCherry and anti-AAF1 antibodies. The free mCherry/full-length mCherry-ATG8d ratio was used as an indicator of autophagic flux. One Ponceau S-stained blot is shown as a loading control. (**c**) Quantification of the free mCherry/full-length mCherry-ATG8d ratio. Bars represent means ± s.e. of three biological replicates, including one replicate from light and two from dark conditions. \*\**P* < 0.01 (two-tailed t test). (**d**) GFP-AAF1-induced symptoms were reduced in *N. benthamiana* leaves subjected to VIGS targeting *RABD2a*. Images were taken 5 days after GFP-AAF1 expression and 20 days after infiltration with VIGS constructs Vi-*rabD1*, Vi-*rabd2a*, Vi-*atg7*, or the Vi-TRV2 empty vector control. (**e**) RT-qPCR analysis of *ATG7*, *RABD1*, and *RABD2a* transcript levels in VIGS plants. Expression levels represent means ± s.e. of three biological replicates, each with three technical replicates, normalized to *Actin* and *PP2A* (*protein phosphatase 2A*). (**f**) Cellular ion leakage, a proxy for cell death, was measured 2 days after infiltration with the GFP-AAF1-expressing construct in VIGS plants. Data represent means ± s.e. of six biological replicates. Different letters in (**e**) and (**f**) indicate significant differences (*P* < 0.05, ANOVA followed by Tukey’s HSD test). Raw data are provided in Data Set S1. (**g**) Comparative LC-MS proteomic analysis of Vi-TRV2 plants expressing GFP-AAF1 (severe cell death symptoms), Vi-*rabd2a* plants expressing GFP-AAF1 (low cell death symptoms), and Vi-TRV2 plants expressing GFP-Frag2 (no cell death symptoms) (four biological replicates each). Differentially abundant proteins are shown as a heat map organized into seven clusters. Cluster members are listed in Table S1, and the complete proteomic dataset is provided in Data Set S3.

Co-expression of GFP-, RFP-, or mCherry-tagged ATG8d with AAF1 in *N. benthamiana* leaf cells revealed partial colocalization in 1–2 µm cytosolic autophagosomes (Fig. S9), although this overlap was substantially less extensive than that observed between AAF1 and the RABD proteins (Figs. 2, S6c, and S9). Overall, while a direct interaction between AAF1 and ATG8d cannot be excluded, our data suggest that AAF1 is not a canonical cargo receptor and that the observed association may be indirect, possibly mediated through associated protein complexes or organelles. A similarly weak association has been reported between an Arabidopsis SAR1 isoform and ATG8e ^45^.

To determine whether AAF1 modulates autophagy, we monitored autophagic flux using mCherry–ATG8d. Upon vacuolar delivery, ATG8 is degraded while mCherry remains stable, allowing flux estimation from the free mCherry/mCherry–ATG8 ratio ^24,95–97^. HA–AAF1 or HA–Frag2 was co-expressed with mCherry–ATG8d in *N. benthamiana* leaves, and flux was measured following either a 6-hour dark or light treatment. In both treatments, HA–AAF1 increased the free mCherry/mCherry–ATG8d ratio, indicating enhanced autophagic flux (Figs. 3b–c). Consistently, GFP–ATG8d assays showed reduced protein levels upon AAF1 expression, supporting increased vacuolar processing (Figs. S10a, b). Free GFP was not reliably detected due to antibody sensitivity. Testing aphid feeding effects on autophagy using GFP–ATG8f was inconclusive (Fig. S10c) and will require additional optimization. Because ubiquitination is closely linked to autophagy and cargo selection ^91,98^, we also examined global ubiquitination. AAF1 expression in *N. benthamiana* resulted in increased ubiquitinated protein species (Fig. S10d).

Together, partial ATG8 association and enhanced autophagic flux suggest that the cell death observed upon high-level expression of AAF1 and RABD1 or RABD2a (Figs. 1a, 2c, 2f, S1, S2a, S4b) may result from excessive autophagic activity. Consistent with this, overexpression of Ypt1 enhances autophagy in yeast ^48^. While autophagy is essential for survival across eukaryotes ^14,98,99^, excessive autophagy can cause irreversible cellular damage and cell death depending on intensity and context ^99–103^. To test the link between AAF1-and RAB-mediated cell death, we used VIGS to silence *RABD1*, *RABD2a*, and *ATG7* in *N. benthamiana*, followed by transient AAF1 expression. AAF1-induced cell death was significantly reduced in Vi-*rabd2a* plants, but not in Vi-*rabd1* or Vi-*atg7* plants (Figs. 3d, S11a). qPCR confirmed strong knockdown of *RABD1* and *RABD2a,* but not *ATG7* (Fig. 3e), consistent with the lack of phenotype in Vi-*atg7* plants. Ion leakage assays confirmed reduced cell death only in Vi-*rabd2a* plants (Fig. 3f), despite comparable AAF1 accumulation (Figs. S11b, c). A similar trend in ion leakage was observed in VIGS plants infiltrated with HA–AAF1 (Fig. S11d). Together, these findings indicate that AAF1-induced symptoms and its function depends, at least in part, on RABD2a.

To further investigate the role of RABD2a in AAF1-induced leaf symptoms, we performed comparative quantitative proteomic analysis by LC–MS on Vi-TRV2 plants expressing GFP–AAF1 (severe cell death), Vi-*rabd2a* plants expressing GFP–AAF1 (reduced cell death), and Vi-TRV2 plants expressing GFP–Frag2 (control; no cell death), with four biological replicates per group (complete dataset in Data Set S3).

Differentially abundant proteins were visualized as a heat map and grouped into seven clusters (Fig. 3g; Table S1). Clusters A, F, and G likely represent AAF1-dependent changes independent of RABD2a, whereas cluster E primarily reflects the effects of *rabd2a* knockdown. In contrast, clusters B and C, and to a lesser extent cluster D, likely represent proteins regulated by AAF1 in a RABD2a-dependent manner. Cluster B contained proteins involved in vesicle trafficking (including a Sec14p-like protein and Rab GTPase homolog A5B) and cytoskeletal organization (including myosin isoforms, a microtubule-associated futsch-like protein, and an ankyrin repeat family protein), all of which accumulated specifically in plants exhibiting severe cell death. Conversely, cluster C contained proteins involved in chorismate biosynthesis, phospholipid metabolism, and vesicle trafficking (SNARE complex components) whose abundance was specifically reduced in these plants. Together, these findings further support a role for RABD2a in mediating AAF1-induced leaf symptoms.

### AAF1 Interacts with PI3PT, an ER-localized Putative Lipid-Binding or Lipid Transfer Protein

The second major interactor identified in the AAF1-binding proteome was XP_016477330 (*LOC107798808*), an uncharacterized protein annotated as a putative phosphatidylglycerol/phosphatidylinositol transfer protein, hereafter referred to as PI3PT. Its Arabidopsis ortholog (*AT3G11780*) is predicted to function in sterol transport and binding. Structural modeling using AlphaFold, Phyre2, I-TASSER, and SMART consistently classified PI3PT as an ML (MD-2-related lipid-recognition) domain-containing protein with strong structural similarity to lipid-binding proteins, including the yeast Niemann–Pick type C protein Npc2 and mammalian MD-1/MD-2 proteins ^104^ (Fig. S12), supporting its predicted role in lipid binding or transfer.

To validate this interaction, GFP-PI3PT and HA-AAF1 were transiently co-expressed in *N. benthamiana*, followed by reciprocal co-immunoprecipitation. GFP-PI3PT efficiently co-immunoprecipitated HA-AAF1 (Figs. 4a, S13a, and S13b). Notably, a fraction of the co-immunoprecipitated AAF1 migrated as SDS-resistant higher-order species distinct from those observed in the AAF1–RABD2a interaction, suggesting that PI3PT may preferentially associate with a specific biochemical form of AAF1. Whether these species reflect post-translationally modified forms of AAF1 remains unknown. Unlike AAF1 or RABD1/RABD2a, prolonged PI3PT overexpression did not induce cell death (Fig. S13c).

**Fig. 4.**
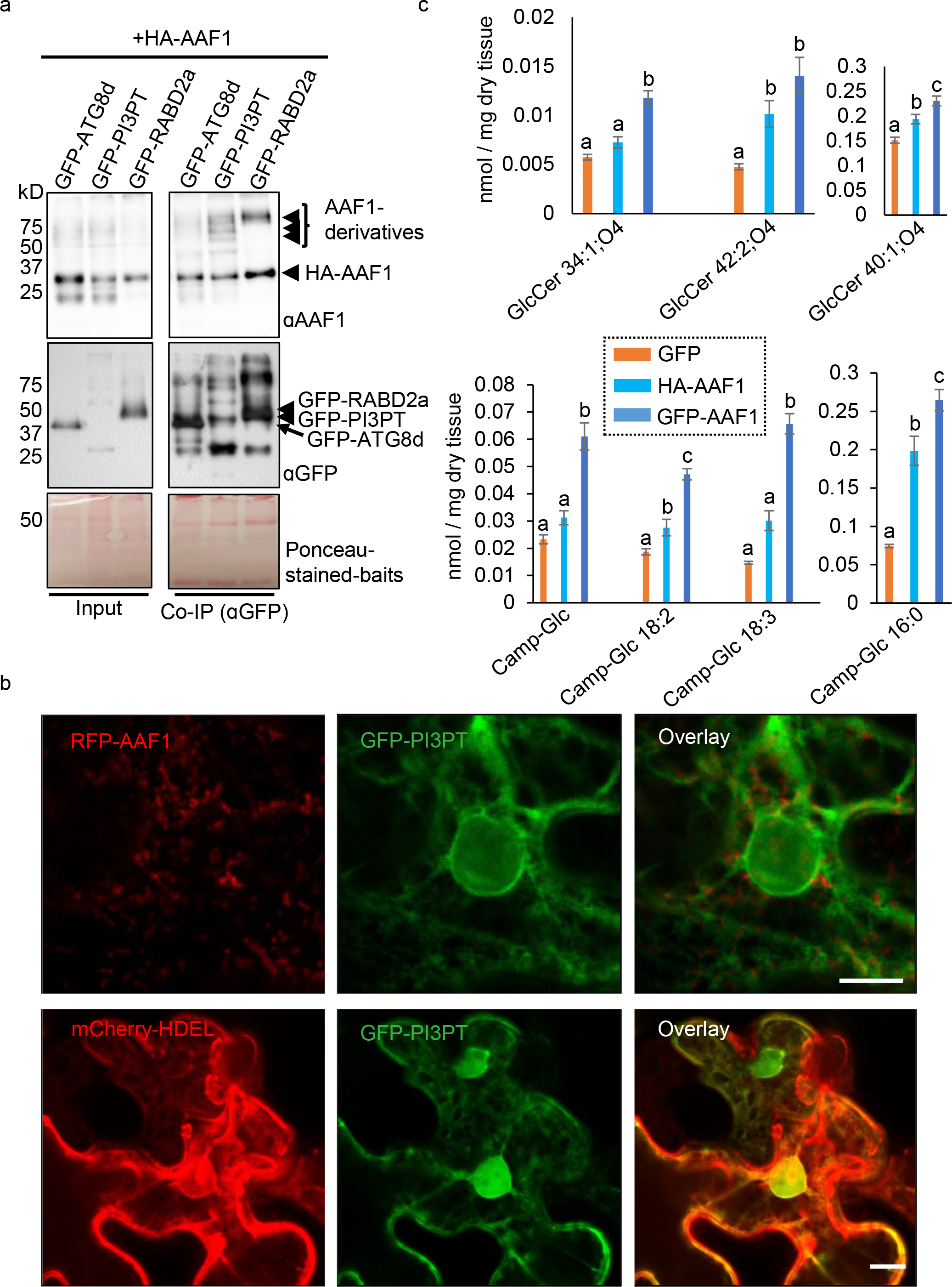
Leaf-expressed AAF1 binds to PI3PT, an uncharacterized ER-localized protein, and promotes the accumulation of specific lipids. (**a**) GFP–PI3PT was used as bait to pull down HA–AAF1. GFP–RABD2a and GFP–ATG8d served as positive control baits. The three upper arrowheads indicate distinct higher-order forms of AAF1, whose immunoblot patterns (replicate experiments shown in Fig. S13) differed depending on the interacting bait protein, RABD2a or PI3PT. Immunoblots of Co-IP eluates and input samples were probed with anti-AAF1 and anti-GFP antibodies. (**b**) Subcellular colocalization of AAF1 with PI3PT in plant cells. Confocal microscopy analysis of *N. benthamiana* leaf epidermal cells coexpressing RFP–AAF1 and GFP–PI3PT showed RFP–AAF1 localized to punctate structures associated with the ER-localized GFP– PI3PT (top panels). ER localization of GFP–PI3PT was confirmed by its colocalization with the ER marker mCherry–HDEL (bottom panels). Scale bar = 10 µm. (**c**) AAF1 expression increases the levels of glucosylceramide (GlcCer) (top bar charts) and campesterol glucoside esters (Camp-Glc) (bottom bar charts) in plants. Total lipids were measured in *N. benthamiana* leaves transiently expressing GFP–AAF1, HA–AAF1, or GFP as a negative control. For each sample, lipid content (nmol lipid per mg dry tissue) was quantified by LC-MS, as described in Methods. Statistical analysis was performed using ANOVA; different letters indicate significant differences (*P* < 0.05, ANOVA followed by Tukey’s HSD test). Raw data are provided in Data Sets S4 and 5.

Subcellular localization revealed that GFP-PI3PT resides in the ER, as confirmed by co-localization with an ER marker (Fig. 4b). RFP-AAF1 puncta were frequently associated with the PI3PT-positive ER network, resembling the organization of pre-autophagosomal structures in mammalian cells ^105^. These observations led us to hypothesize that PI3PT may either (i) recognize PI3P or other lipids on the ER or ER-contact membranes to recruit the AAF1–RABD2a complex or (ii) facilitate lipid transfer between the ER and adjacent membranes. However, its precise function remains unknown.

Because PI3PT is predicted to function in lipid binding or transfer, we examined whether AAF1 expression alters lipid homeostasis by performing lipidomic mass spectrometry on *N. benthamiana* leaves expressing high or low levels of AAF1, using GFP-expressing leaves as controls. AAF1 expression significantly increased several lipid classes (Figs. 4c, S14a, b; Supplementary Data Sets S4 and S5), including glucose ceramides, sterol esters, glucosylated sterols, and triacylglycerols. These lipid groups have established roles in autophagy and lipid droplet (LD) biogenesis ^33,99,106–110^.

Because sterol esters and triacylglycerols are major LD constituents, we examined the spatial relationship between LDs and AAF1 puncta by staining leaves expressing RFP-AAF1 with BODIPY. Confocal imaging revealed that LDs frequently surrounded and accumulated around AAF1-positive puncta (Fig. S14c). Interestingly, PEXRD54, an effector from *P. infestans* that shares some functional similarities with AAF1, has also been reported to facilitate starvation-induced autophagy by recruiting LDs to the fungal infection site ^59^. Whether AAF1-mediated lipid changes promote autophagosome biogenesis or instead reflect cell death-or lipophagy-associated lipid remodeling remains to be determined.

### AAF1 and RABD2a Interact on ER-Associated Mobile Compartments Involved in Autophagosome Formation or Trafficking

To validate the interaction between AAF1 and RABD2a independently of Co-IP, we used a bimolecular fluorescence complementation (BiFC) system ^111,112^. AAF1 or Frag3 (a truncated version of AAF1 used as a negative control, lacking the RBD and an additional upstream region, making it shorter than Frag2 shown in Fig. 2) was fused to the N-terminal half of YFP (nYFP), whereas RABD2a or RABD1 was fused to the C-terminal half of YFP (cYFP). mCherry– GUS was expressed between the two cassettes as a transformation marker. We observed highly mobile compartments that continuously changed shape while moving throughout the cell, formed only in combinations containing full-length AAF1 and either RABD protein (Fig. S15a; Videos 1a and b). Cytosolic GUS–mCherry (red fluorescence) was present at very low levels and was not detectable above background.

Compartments marked by the AAF1–RABD1 complex were less abundant than those marked by AAF1–RABD2a, whereas no mobile compartments were observed with Frag3, indicating that formation of these compartments requires full-length AAF1 (Fig. S15a). Typically, a single compartment measuring 2–8 µm was observed per cell. Co-expression with the nuclear marker GTE4-RFP confirmed that these mobile compartments were distinct from the nucleus (Fig. 5a; Video 2). Unlike the TGN/EE puncta ^62–64^ marked by the colocalization of RABD and AAF1 proteins expressed from pEAQ vectors (Fig. 2), the AAF1–RABD BiFC compartments consistently appeared as mobile puncta that developed into larger compartments in some cells. This difference could reflect the lower and potentially more biologically relevant protein levels produced by the BiFC vectors, which may permit more natural subcellular trafficking, or altered membrane dynamics associated with BiFC complex formation. Regardless of the underlying mechanism, the BiFC system enabled visualization and tracking of AAF1– RABD-associated compartments.

**Fig. 5.**
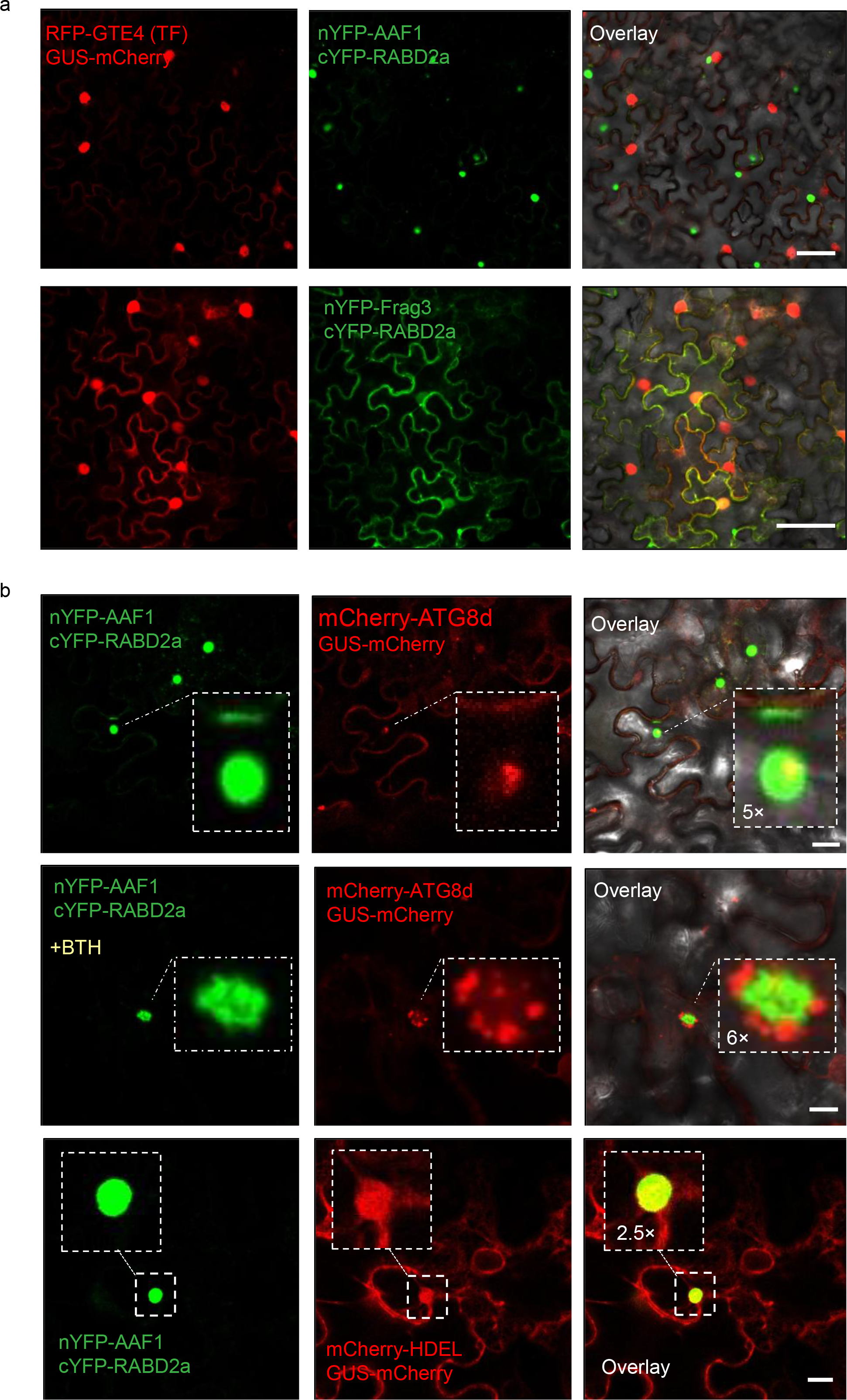
Interacting AAF1 and RABD2a form previously uncharacterized mobile compartments with overlapping ER composition that generate and/or traffic autophagosomes (Videos 1–6 support this figure). (**a**) Images captured by sequential confocal scanning in the top row show compartments marked by the complemented interaction of nYFP–AAF1 and cYFP–RABD2a (green fluorescence) and nuclei marked by RFP–GTE4, a transcription factor used as a nuclear marker (red fluorescence), following co-expression from independent plasmids in *N. benthamiana*, showing the relative size and position of the AAF1–RABD2a–containing compartments relative to nuclei. The second row shows that co-expression of nYFP–Frag3 (a truncated AAF1 lacking the Rab-binding domain; negative control) and cYFP–RABD2a does not result in detectable BiFC-labeled mobile compartments. Scale bars = 50 μm. (**b**) The top two rows show compartments marked by the complemented interaction of nYFP–AAF1 and cYFP–RABD2a (green fluorescence), autophagosomes marked by mCherry–ATG8d (red fluorescence), and their overlap (yellow), following co-expression from independent plasmids (corresponding images from a different leaf area are shown in Fig. S15b). The second row shows a leaf area sprayed with 100 µM BTH, which induces autophagosome formation. The third row shows mCherry–HDEL (red fluorescence) co-expressed from a separate plasmid together with nYFP–AAF1/cYFP–RABD2a to show the localization and overlap of the compartments (green fluorescence) relative to the ER (red fluorescence) and their overlap (yellow). Insets in the first, second, and third rows show 5×, 6×, and 2.5× magnified views of the indicated areas, respectively. Scale bars = 10 μm. Except for the third row in (**b**), the overlay panels include bright-field images. GUS–mCherry (red fluorescence) was co-expressed from the same T-DNA as nYFP– AAF1 and cYFP–RABD2a and served as a cytosolic transformation marker in all images. Because GUS–mCherry expression from the BiFC vector was low and negligible (Fig. S15a), we did not expect it to interfere with the detection of autophagosomes (mCherry–ATG8d) or the ER (mCherry–HDEL).

To determine whether these compartments are associated with autophagosomes, we co-expressed mCherry-ATG8d together with the AAF1–RABD2a BiFC constructs. Confocal imaging and time-lapse microscopy showed autophagosomes forming on, or being trafficked by, the mobile compartments (Fig. 5b, top and middle rows; Videos 3a–c and 4a–c). Although mCherry-ATG8d fluorescence was faint and diffuse within the compartment, it was strongly enriched in autophagosomes. Following induction of autophagy with BTH (benzo[1,2,3]thiadiazole-7-carbothioic acid S-methyl ester), the compartments became less mobile while accumulating increased numbers of autophagosomes (Fig. 5b, middle row). These observations support the hypothesis that the AAF1–RABD2a compartment functions as a platform for autophagosome formation and/or trafficking. To examine the relationship between the compartment and the ER, we coexpressed mCherry-HDEL with the AAF1– RABD2a BiFC constructs. Time-lapse imaging revealed the presence of the ER marker within the compartment and continuous exchange of ER membrane between the compartment and the surrounding ER network (Fig. 5b, bottom row; Videos 5a–c and 6a–c).

Overall, the high motility of these AAF1–RABD compartments suggests two possible identities. First, they may represent enlarged early, late, or recycling endosomes. Endosomes can exhibit elongated or vesicular morphologies, are highly mobile, and can reach several microns in size. Pathological or environmental disruption of endosomal trafficking can cause endosomes to enlarge to up to 10 µm ^113,114^. In certain cell types, large endocytic vesicles, or macropinosomes, which engage in crosstalk with autophagy during starvation and require RAB cascade during maturation, can reach similar sizes ^115–117^. Recent studies in mammalian cells have shown that phagophores can arise from recycling endosomes ^81–83^. Specifically, the mammalian PI3P-binding protein WIPI2, an ATG18 homolog, binds PI3P on recycling endosomes associated with the RAB GTPase RAB11A, thereby recruiting the ATG5–ATG12– ATG16L1 complex and initiating autophagosome formation on endosomal membranes ^82^. Endosomal involvement in autophagy is not limited to autophagosome formation. Independent studies in both animals and plants have shown that endosomes and multivesicular bodies (endolytic compartments derived from the cell surface) can fuse with autophagosomes to form highly mobile and relatively large (∼2 µm) amphisomes, which subsequently traffic to lysosomes or vacuoles for cargo degradation ^118,119^.

Alternatively, the AAF1–RABD-containing compartments may represent ER– endosome contact sites (EERCSs), which have been proposed to form membraneless compartments through liquid–liquid phase separation. Such biomolecular condensates can reach several microns in size and are highly mobile. EERCSs increase as endosomes traffic and mature ^120^. A recent study in mammalian cells further suggested that autophagosomes can form on endosomes through molecular machinery associated with EERCSs and that the dynamic mobilization of EERCSs creates a confined cytoplasmic environment that promotes Ca²⁺ accumulation and liquid–liquid phase separation, thereby establishing platforms for de novo autophagosome biogenesis ^80^.

Taken together, these findings suggest that AAF1 and RABD may function in endosomal trafficking and autophagy. However, the precise nature of the mobile AAF1– RABD compartments observed in our study remains to be determined.

### AAF1 Expression Promotes Plant Growth via an Autophagy-Dependent Mechanism

Because enhanced autophagy has been associated with improved nutrient recycling and plant growth ^15–17,19,26,27^, we investigated whether constitutive AAF1 expression promotes plant growth in an autophagy-dependent manner. AAF1-HA was expressed at low to moderate levels in Arabidopsis Col-0 and the autophagy-deficient *atg5* mutant, both carrying an ectopic copy of the Arabidopsis ATG8a gene fused to GFP under the control of the ubiquitin promoter. Homozygous transgenic lines were selected for subsequent analyses. Compared with wild-type Col-0 plants, AAF1-expressing plants exhibited significant increases in seedling fresh weight, seedling surface area at both early and late developmental stages, mature rosette size, seed size, and dry biomass (Figs. 6a–h and S16). In contrast, these growth-promoting effects were abolished in the *atg5* background, indicating that AAF1-dependent growth requires a functional autophagy pathway.

**Fig. 6.**
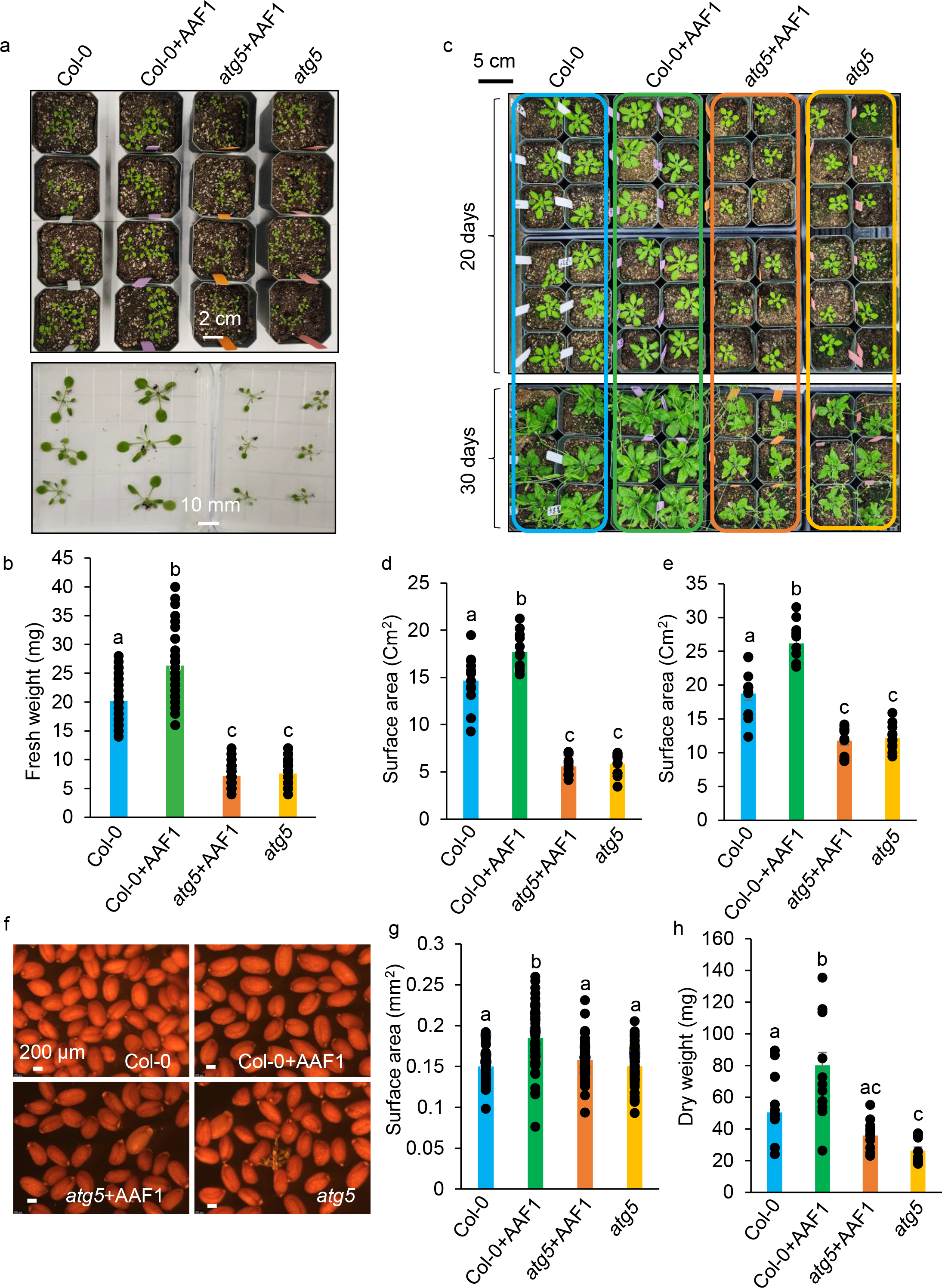
AAF1 enhances plant growth, biomass, and aphid fecundity in Arabidopsis. Wild-type Col-0 and *atg5* Arabidopsis plants, both already expressing At-Ubi:ATG8a, were transformed with AAF1-HA(3×) and advanced to the T3 generation. Transformed plants expressing were grown alongside their corresponding untransformed wild-type and *atg5* mutant plants until maturity. (**a**–**b**) Fresh weight of 2-week-old seedlings was measured after removal from the soil. Data represent the mean ± s.e. (N = 55 for Col-0, N = 42 for Col-0+AAF1, N = 76 for *atg5*+AAF1, and N = 74 for *atg5*). (**c**–**e**) Rosette surface areas of 20-day-old and 30-day-old seedlings from the same growth experiment were measured using Fiji after capturing images of each plant at the same scale. Data represent the mean ± s.e. (N = 12 for all genotypes in both cases). (**f**–**g**) Seed sizes were measured using Fiji after capturing images of seeds from each plant at the same scale using a Leica M205 stereoscope. Data represent the mean ± s.e. (N = 60 for all genotypes). (**h**) Dry weight of mature plants (35 days old) from a different growth experiment was measured (N = 11 for Col-0 and *atg5*, and N = 12 for Col-0+AAF1 and *atg5*+AAF1). Statistical analysis was performed using ANOVA; different letters indicate significant differences (*P* < 0.001, ANOVA followed by Tukey’s HSD test).

Immunoblot analysis revealed enhanced autophagic flux in AAF1-expressing wild-type plants, whereas autophagic flux remained impaired in *atg5* plants regardless of AAF1 expression (Fig. 7a). Surprisingly, neither GFP–ATG8a nor free GFP was detected in *atg5* mutant plants with the Roche anti-GFP antibody, regardless of AAF1 expression, even though both genotypes expressed GFP–ATG8a and showed elevated GFP–ATG8a accumulation when probed with a different anti-GFP antibody (BioLegend). Consistent with autophagy-dependent turnover, AAF1 accumulated to higher levels in the *atg5* mutant and decreased under dark treatment, suggesting that AAF1 itself undergoes autophagic degradation (Fig. 7a; Fig. S16c).

**Fig. 7.**
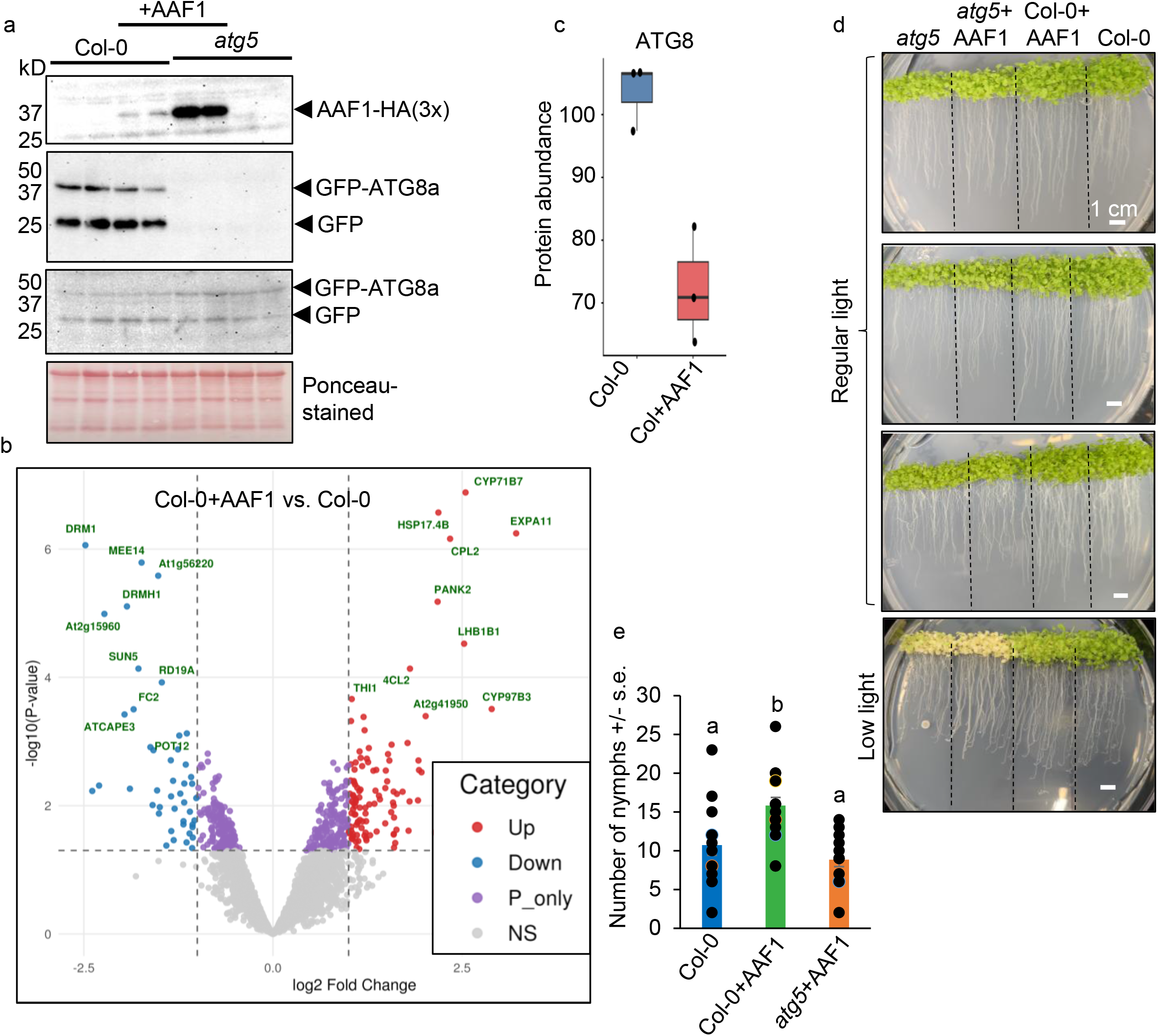
Autophagy-inducing activity of AAF1 in Arabidopsis is associated with additional phenotypic traits. (**a**) Immunoblot of total proteins from pooled seedlings of each of the four Arabidopsis genotypes harvested from the same plant batches shown in Fig. 6a. Blots were probed with anti-AAF1 and two anti-GFP antibodies: Roche (Cat. No. 11814460001; top) and BioLegend (Cat. No. 902605; bottom). (**b**) Comparative LC-MS proteomic analysis of pooled seedlings harvested from the same plant batches shown in Fig. 6a and used for the immunoblot analysis in Fig. 7a. Volcano plot showing differential protein abundance between Col-0+AAF1 and Col-0 (three biological replicates per genotype). P_ only, *P*-value cutoff; ns, not significant. Differentially abundant proteins are listed in Table S2, and the complete proteomic dataset is provided in Data Set S6. (**c**) Comparison of ATG8 protein abundance between Col-0 and Col-0+AAF1 based on the quantitative proteomic analysis. (**d**) Two-week-old seedlings of the four genotypes were grown vertically on ½ MS medium without sucrose to monitor root growth. The three upper plates represent independent experiments; one experiment (bottom) was horizontally maintained under low light for an additional 7 days. (**e**) Individual one-day-old *M. persicae* nymphs were caged on a single rosette leaf of plants from three of the four genotypes, and progeny were counted 7 days later. Data represent means ± s.e. (Col-0, N = 15; Col-0+AAF1, N = 19; *atg5*+AAF1, N = 13). Different letters indicate significant differences (*P* < 0.01, one-way ANOVA followed by Tukey’s HSD test).

To identify downstream changes associated with AAF1 expression, we performed quantitative proteomic analysis using pooled seedlings harvested from the same plant batches shown in Fig. 6a and used for the immunoblot analysis in Fig. 7a. Three biological replicates (10 seedlings per pool) were analyzed for each genotype. Results are presented in Figs. 7b and S17, Table S2, and Data Set S6. In the wild-type background, AAF1 increased the abundance of proteins associated with cell expansion and growth (EXPA11, a member of the alpha-expansin family), cell wall biosynthesis (4CL2, an isoform of 4-coumarate ligase), photosynthesis (LHB1b1, a Photosystem II type I chlorophyll a/b-binding protein), growth regulation (CPL2, carboxyl-terminal domain [CTD] phosphatase-like 2) ^121^, and thiamine biosynthesis (THI1, thiamine thiazole synthase 1), which has also been implicated in plant growth and stress resilience ^122^ (Fig. 7b, Table S2). These changes were largely absent in the *atg5* background (Fig. S17; Table S2). Proteins associated with dormancy and DNA methylation, including DRM1 (Domains Rearranged Methylase 1), as well as DRMH1 and DRMH3, members of the dormancy/auxin-associated protein family, were reduced in AAF1-expressing wild-type plants (Fig. 7b, Table S2). The proteomic analysis also revealed reduced ATG8 abundance, consistent with increased autophagic flux (Fig. 7c).

Wild-type plants expressing AAF1 also developed significantly longer roots than the other genotypes (Fig. 7d), suggesting that this enhancement may underlie, or be associated with, the overall growth-promoting effect of AAF1. The importance of autophagy in root development and growth has recently been demonstrated ^123^. Furthermore, treatment of roots with concanamycin A, with or without BTH, resulted in greater accumulation of GFP-ATG8a-labeled autophagosomes and autophagic bodies in AAF1-expressing roots than in wild-type controls (Fig. S18), providing additional evidence for enhanced autophagy.

Finally, aphid fecundity was significantly higher on AAF1-expressing Col-0 plants than on Col-0 wild-type control and the *atg5* mutant expressing AAF1 (Fig. 7e). An independent experiment that included all four genotypes produced the same trend, although the differences were not statistically significant because of the reduced number of biological replicates (Fig. S19). Collectively, these findings demonstrate that AAF1 enhances plant growth through an autophagy-dependent mechanism while increasing host susceptibility to aphid colonization.

## Conclusions

Together, our findings show that AAF1 interacts and colocalizes with plant RABD proteins on TGN/EE puncta, which have previously been implicated in autophagosome formation. We further show that RABD2a exhibits greater specificity than RABD1 in mediating AAF1 function. We identified highly mobile compartments marked by the AAF1–RABD complex and demonstrated that these compartments are associated with the ER and traffic with and/or potentially generate autophagosomes. We also found that AAF1 binds to an uncharacterized ER-localized lipid-binding protein, suggesting a possible role in mobilizing ER-derived lipids required for autophagosome biogenesis and expansion. Based on our data, we identified functional parallels with two recent models from mammalian cells. The first model indicates that phagophores can initiate and expand on recycling endosomes (Fig. 8a, adapted from ^82^) through the coordinated action of PI3P-binding proteins and endosomal Rab GTPases, which recruit autophagosome initiation complexes to endosomal membranes ^81–83^. The second proposes that membraneless biomolecular condensates at ER–endosome contact sites (EERCSs), formed through liquid–liquid phase separation, are highly mobile and provide an environment conducive to autophagosome formation on endosomes ^80^ (Fig 8b). We observed functional parallels in plants and propose that the PI3PT–AAF1– RABD2a module forms a bridge at ER–endosome contact sites. This complex may facilitate the transfer of ER-derived membranes and lipids to expanding autophagosomes or contribute to the formation of compartments with an endosomal or EERCS-like identity (Fig. 8c). We further propose that aphids exploit this process to accelerate starvation-induced autophagosome formation and nutrient recycling at feeding sites, thereby creating a more favorable environment for their survival. Finally, we show that ectopic expression of AAF1 enhances plant growth and biomass in an autophagy-dependent manner.

**Fig. 8.**
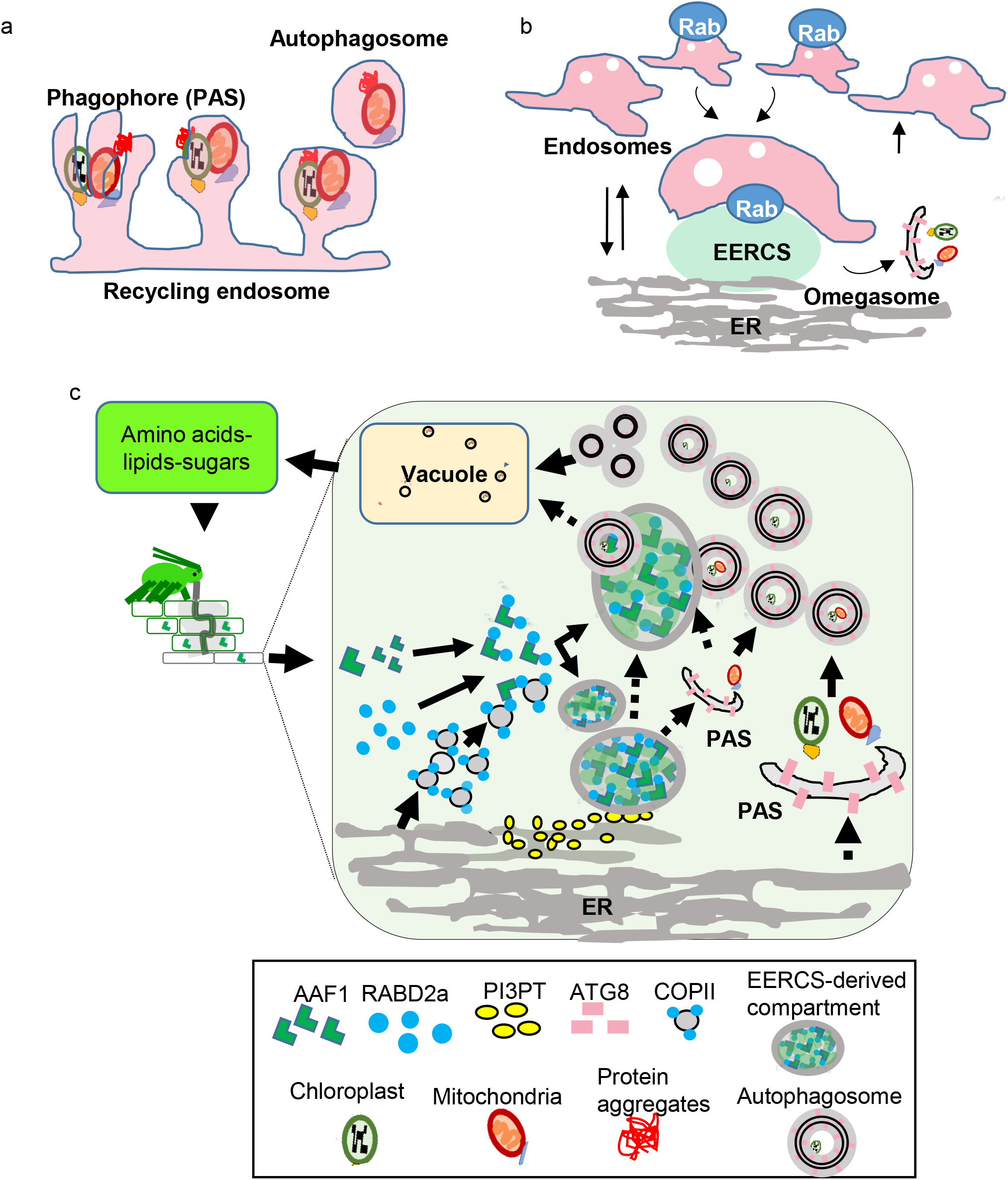
Schematic representation of AAF1 function in plants. (**a**) A model for phagophore formation on recycling endosomes in mammalian cells, adapted from ^82^, and (**b**) a follow-up model for phagophore formation at endosome–ER contact sites (EERCSs), adapted from ^80^, showing similarities to our proposed model (**c**) for autophagosome formation. (**c**) Aphid-secreted AAF1 enters the plant cell and binds RABD2a on an EERCS-derived compartment that acts as a platform for the generation and/or trafficking of autophagosomes before their delivery to vacuoles. AAF1 also binds an ER-localized protein, PI3PT, which contains putative lipid-binding domains but whose function remains undetermined. We propose that the RABD2a–AAF1–PI3PT complex is at the nexus of autophagosome formation at EERCSs. These interactions are proposed to enhance vesicle and/or lipid mobilization, supplying membrane material for autophagosome formation, which may consequently promote nutrient recycling and thereby benefit aphid feeding and performance.

## Methods

### Plants, Insects, and Growth Conditions

Wild-type *Arabidopsis thaliana* Columbia-0 (Col-0) and the *atg5* mutant, both expressing At-*ATG8a* under the ubiquitin promoter, were obtained from the Arabidopsis Biological Resource Center (https://abrc.osu.edu/) and Dr. Céline Masclaux-Daubresse (Université Paris-Saclay, France), respectively.

Wild-type and *asat2-1* mutant *Nicotiana benthamiana* ^124^, as well as wild-type and *atg5* mutant Arabidopsis, were grown in Cornell Mix (by weight: 56% peat moss, 35% vermiculite, 4% lime, 4% Osmocote slow-release fertilizer [Scotts, Marysville, OH], and 1% Unimix [Scotts]) in Conviron (Winnipeg, Canada) growth chambers under a 16:8 h photoperiod, 180 µmol m⁻² s⁻¹ photosynthetic photon flux density, and a constant 23 °C temperature. A genome-sequenced red strain of *Myzus persicae* ^125,126^, maintained on *Nicotiana tabacum* under similar light conditions, was used for all experiments.

### Preparation of Aphid Salivary Proteome

Approximately 200 mg mixed-instar *M. persicae* were transferred into modified small cups with mesh bottoms and tops sealed with two layers of Parafilm (Fig. S20). An artificial diet consisting of 200 mM sucrose was placed between the Parafilm layers. After two days of feeding, the diet, now mixed with aphid saliva, was collected and pooled from ten cups. The solution was then concentrated and further processed to remove sucrose using Centricon spin columns (Millipore Sigma) with a 3 kDa molecular weight cutoff, and subsequently used for LC-MS/MS analysis.

### Tandem Mass Spectrometry LC-MS/MS Analysis of Aphid Salivary Proteins and Co-IP Eluates

Protein samples were reduced in 5 mM TCEP at 56°C for 45 min, alkylated in 10 mM IAA at 23 °C for 30 min, and digested by adding 1 µg trypsin (Trypsin Gold, Mass Spectrometry Grade, Promega) at 40 °C overnight. LC-MS analysis of peptides was performed by using a Dionex Ultimate 3000 Series LC system controlled by Chromeleon software (Thermo Fisher Scientific) coupled to a heated electrospray ionization source (HESI-II) on a Q-Exactive mass spectrometer controlled by Xcalibur software (Thermo Fisher Scientific). Peptide mixtures were separated on a C18 column 150 x 2.1 mm, 1.7µm, 100 Å (Aeris PEPTIDE XB, Phenomenex) at 50 °C with a multi-stepped two-buffer gradient elution (buffer A, 0.1% formic acid; buffer B, acetonitrile,0.1% formic acid) and a flow rate of 250 µl/min. The mass spectrometer was externally calibrated, and the HESI source was set to spray voltage +3500 V, probe heater temperature 400 °C, and sheath gas and auxiliary gas flows 50 and 10 arbitrary units, respectively. The source optic was set to capillary temperature 350 °C and S-Lens RF Level 70. Data were acquired under the Scan Mode FTMS + p Full MS/ Data Dependent MS2 (TopN). The MS1 scans were conducted at resolution 70,000, default charge state 2, AGC target 5e6, maximum IT 50 s, scan range 350 to 1000 m/z, chrom. peak width (FWHM) 4 s and the dd-MS² at resolution 17,500, AGC target 1e5, maximum IT 50 s, loop count 10, TopN 10, isolation window 2.0 m/z, stepped (N)CE 15, 30 (%), with filter settings of minimum AGC target 1e3, intensity threshold 2e4, charge exclusion of unassigned, 1, 6-8, >8, exclude isotopes, on the dynamic exclusion mode at mass tolerance low = ±10, exclusion duration 15 s, exclusion after 1 time, and filter intensity threshold of minimum 10,000. Protein identification and grouping: The LC-MS/MS raw data were processed, and proteins identified using Proteome Discoverer 2.2 (Thermo Fisher Scientific). The processing workflow consisted of Spectrum Selector, Sequest HT, and fixed value peptide-to-spectrum match validator nodes. The Consensus workflow consisted of Peptide and Protein Filter, Protein Scorer, Protein false discovery rate Validator, and Protein Grouping nodes. The search parameters were set to the precursor mass tolerance of 0.006 Da, fragment mass tolerance of 0.02 Da, up to three missed cleavages allowed, minimal peptide length of six amino acids, static modification: carbamidomethylation [C], dynamic modifications: oxidation [M]; deamidation [Q]; acetylation [S/T/K/H/Y/N-term], Max Delta Cn of 0.01, target false discovery rate of 0.01, search database of NCBI Refseq *N. tabacum* and *N. benthamiana*, protein grouping of Gene Ontology database (Thermo Fisher Scientific). For the Co-IP experiments, the sorted list of identified proteins, their corresponding peptides, and detailed information have been provided in Data Set S2.

### Tandem Mass Spectrometry LC-MS/MS Analysis of Proteins from Arabidopsis and *N. benthamiana* Leaf Tissues

Arabidopsis and tobacco total proteins were extracted using the protocol described ^127^. Briefly, ∼100 µg of protein were used for downstream processing. DTT and IAA were used for reduction and alkylation, respectively. Subsequently, trypsin/LysC digestion was performed. The resulting peptides were desalted using Thermo Fisher C18 desalting columns. Mass spectrometer settings were kept identical to those described in ^127^, with the only modification being the injection of ∼300 ng of peptides per sample. Instrument performance was assessed using a HeLa digest run before and after sample acquisition. Raw (.d) files were analyzed using Spectronaut v20 in direct DIA mode. For Arabidopsis, the Arabidopsis thaliana FASTA proteome was used to generate the spectral library, and for tobacco (*Nicotiana sylvestris*), the corresponding FASTA file was used. All settings were kept at default values. Protein abundance matrices generated from mass spectrometry-based proteomics experiments were preprocessed using custom scripts prior to statistical analysis. Missing values were addressed using a hybrid imputation strategy combining k-nearest neighbors (KNN) and minimum detection (MinDet) methods to account for both missing-at-random (MAR) and missing-not-at-random (MNAR) mechanisms. Within each experimental condition, protein intensity values equal to zero were treated as missing observations and converted to NA values. For each protein, the proportion of observed measurements within each condition was calculated. Proteins with at least 60% observed values were classified as MAR candidates and imputed using KNN, whereas proteins with fewer than 60% observed values were considered likely MNAR and imputed using the MinDet approach. KNN imputation was performed using the scikit-learn implementation of the KNNImputer algorithm with k = 3 nearest neighbors. Missing values were estimated from proteins with similar abundance profiles, thereby preserving local data structure and biological relationships. Proteins classified as MNAR were imputed using the MinDet approach: for each condition, the minimum detectable protein intensity was identified after excluding missing and zero values, and missing observations were replaced with 90% of this minimum detected abundance. Following condition-specific imputation, a final quality-control step was applied across the complete dataset. Any remaining missing values were replaced using a global MinDet estimate derived from the lowest detected abundance across all samples, resulting in a complete quantitative matrix for downstream analyses. The imputed abundance matrix was imported into proteoDA ^128^ for normalization and quality assessment. Protein abundance data were then imported into R and organized using the DAList framework, with sample metadata linked through unique sample identifiers. Different normalization methods were evaluated to identify the approach that best reduced technical variation while preserving biological differences. Performance was assessed using principal component analysis (PCA), hierarchical clustering, sample-to-sample distance metrics, abundance distributions, and variance estimates to evaluate both inter- and intra-group relationships. Based on these quality-control assessments, VSN normalization was selected for downstream differential abundance analysis because it provided the most consistent variance stabilization and clustering of biological replicates. Differential protein abundance analysis was performed using the limma package ^129^, which employs linear modeling and empirical Bayes variance moderation to provide robust statistical inference, particularly for studies with relatively small sample sizes. Normalized protein abundance values were organized according to experimental groups, with sample assignments inferred from sample naming conventions and encoded as categorical factors. A design matrix without an intercept was constructed to estimate group-specific abundance levels independently, and differential abundance between experimental groups was evaluated using user-defined contrast matrices. Differentially abundant proteins were identified using a nominal p-value threshold of ≤ 0.05 and an absolute log2 fold-change threshold of 1. Corresponding heat maps and volcano plots were generated. Raw quantitative proteomics data have been deposited in the repository under accession ID MSV000101945. The complete proteomic dataset is provided in Data Sets S3, and S6.

### Cloning Procedures and Plasmid Constructs

All primers used in this study are listed in Supplementary Data Set S7. RNA was extracted from aphid bodies using the Qiagen RNeasy Kit (Cat #: 74904). Complementary DNA (cDNA) was synthesized from the extracted RNA using random hexamers and the Applied Biosystems™ High-Capacity cDNA Reverse Transcription Kit. For *in planta* protein expression, gene-specific primers (Supplemental Data Set S7) were designed to amplify the coding regions of selected aphid salivary protein genes, including *AAF1*, based on sequences in the aphid database (https://bipaa.genouest.org/is/aphidbase/), excluding the signal peptides. Amplicons were generated using Phusion proofreading polymerase (New England Biolabs, USA) and subsequently BP-cloned into the pDONR207 Gateway entry vector using the Gateway™ BP Clonase™ II Enzyme Mix (Invitrogen, Carlsbad, CA, USA). Genes encoding ATG8c, ATG8d, ATG8f, RABD1, RABD2a, and PI3PT proteins were amplified from cDNA synthesized from *N. tabacum* RNA extracts, and then cloned into the pDONR207 entry vector (Addgene). Stop codons were either retained or removed, depending on the tag position in the destination vectors. Destination constructs expressing GFP-AAF1, GFP-tagged AAF1 fragments (Frags 1–3), GFP-ATG8c, GFP-ATG8d, GFP-ATG8f, GFP-RABD1, GFP-RABD2a, and GFP-PI3PT were generated via Gateway™ LR recombination (Invitrogen) of the corresponding pDONR207 entry clones into the pEAQ-EGgW destination vector ^65^. The construct expressing Catalase 1-GFP was generated by LR recombination into the pEAQ-gwEG vector; RFP-AAF1, RFP-RABD1, and RFP-RABD2a were cloned into pEAQ-TRgw; and AAF1-RFP was cloned into the pEAQ-gwTRH destination vector ^65^. The corresponding genes were first cloned into pDONR207 via BP recombination. The gene encoding *Flag-AAF1* was obtained through gene synthesis from Twist Bioscience (https://www.twistbioscience.com/) and then cloned into the pDONR207 entry vector before LR cloning into pEAQ-gWEG destination vector ^65^. The destination construct expressing mCherry-ATG8d, AAF1-Flag, and AAF1-HA(3x) were generated by Gateway™ LR recombination (Invitrogen) of the corresponding pDONR207 entry clones into the pK7 m34GW destination vector ^130^.

Destination constructs expressing HA-AAF1 and HA-AAF1 fragments (Frags) 1–3, were generated by cloning AAF1 or its truncated versions into pDONR P2R–P3 (Invitrogen) entry vector, followed by recombination into the pK7 m34GW destination vector ^130^. To generate BiFC double-expression vectors, AAF1 or its truncated version, Frag3, together with either RABD1 or RABD2a, were cloned into the pDONR P2R–P3 (Invitrogen) entry vector and subsequently recombined into a modified pCAMBIA2300 double-expression vector ^111^. YFP complementation was visualized in transformed cells exhibiting fluorescence from the transformation marker GUS–mCherry using confocal sequential scanning. The gene from *N. tabacum* encoding XP_016473241.1, orthologous to the Arabidopsis GTE4 transcription factor (AT1G06230), was amplified using the primers described in Data Set S7 and cloned into pDONR207 before being transferred into pEAQ-TRgw. Vectors used for VIGS, Vi-*AAF1*-1, Vi-*AAF*1-2, Vi-*atg7*, Vi-*rabD1*, and Vi-*rabD2a*, were generated by amplifying 200–300 bp segments of the corresponding genes, cloning them into the pDONR207 entry vector, and subsequently transferring them into Gateway-compatible TRV2 (pTRV2) destination vectors ^131^. Modified versions of pEAQ vectors were kindly provided by Dr. Corinne Schmitt-Keichinger (Université de Strasbourg, France) ^65^. Vectors for expression of GFP and mCherry–HDEL were kindly provided by Dr. Maria Harrison ^130^.

### Protein Transient Expression, Virus-Induced Gene Silencing (VIGS)

Transient protein expression in *N. benthamiana* was performed as previously described ^132^. Briefly, destination vectors were transformed into *Agrobacterium tumefaciens* strain GV3101 and cultured in LB medium containing either 50 mg l⁻¹ kanamycin or 50 mg l⁻¹ streptomycin (depending on the vector), along with 50 mg l⁻¹ rifampicin, for one day. Cultures were centrifuged at 2,000 × g for 10 minutes, and the pellets were resuspended in infiltration buffer (10 mM MES, 10 mM MgCl₂, and 200 μM acetosyringone). The suspensions were adjusted to an OD₆₀₀ of 0.5 and infiltrated into young but fully expanded *N. benthamiana* leaves from 3-to 4-week-old plants using a needleless syringe. The same procedure was used for VIGS, except that 10-day-old seedlings were used for infiltration with VIGs constructs and plants were used for further experiments two weeks post-infiltration.

### Arabidopsis Transformation and Growth on Plates

Arabidopsis floral dipping was performed as previously described ^133^. For growth on plates, seeds were surface-sterilized with 50% bleach and 0.1% Tween for 10 minutes, followed by five washes with sterile distilled water. Sterilized seeds were grown vertically on ½ Murashige and Skoog (MS) medium containing 0.5% phytoagar without added sucrose.

### Leaf Processing, Imaging, and Image Analysis

Hydrogen peroxide (H₂O₂) levels in leaves transiently expressing proteins were measured using the Amplex® Red Hydrogen Peroxide/Peroxidase Assay Kit (Invitrogen), following the manufacturer’s instructions. Subcellular localization of fluorescent proteins was visualized using a Zeiss LSM 800 laser scanning confocal microscope equipped with argon and DSS561 diode lasers. GFP fluorescence was excited at 488 nm and detected between 497–526 nm. RFP and mCherry fluorescence were excited at 561 nm and detected between 600–638 nm. YFP was excited at 514 nm and emitted fluorescence was collected from 529 to 549 nm. For simultaneous imaging of YFP and mCherry/RFP fluorescence during BiFC, sequential scanning was applied. Differential interference contrast (DIC) images were collected simultaneously with the fluorescence images. To visualize lipid droplets, 10 µM BODIPY™ 493/503 (4,4-Difluoro-1,3,5,7,8-Pentamethyl-4-Bora-3a,4a-Diaza-s-Indacene) (Invitrogen) was infiltrated into *N. benthamiana* leaves two days after infiltration with vectors for transient expression of RFP-AAF1.

### Aphid Bioassays

For the fecundity assay on AAF1-expressing leaves, one-day-old nymphs were first reared on detached tobacco leaves placed on agar medium before being transferred to leaves of the *N. benthamiana asat2-1* mutant that had been infiltrated with *AAF1* constructs the previous day. One nymph per leaf was confined using polyester mesh fruit netting bags (10 × 15 cm; Amazon), which were manually sealed and secured at the opening with glued cotton balls after placement. After 7 days, the mesh bags containing the leaves were detached from the plants, and the number of nymphs produced was counted. Other fecundity bioassays (Figs. S4 C and D) were performed using custom-made cages constructed from the cut tops of 50 mL Falcon tubes. The tubes were capped with modified lids containing mesh inserts, and the cages were secured onto leaves using pieces of Styrofoam and nails fixed into the tubes. The survivorship bioassay on VIGS-*AAF1* plants (Fig. 1E) was performed as previously described ^70^. For the fecundity assay on Arabidopsis plants, one-day-old nymphs were first reared as described above before being transferred to attached Arabidopsis rosette leaves and confined using custom-made cages constructed from small cups with modified bottoms containing mesh inserts and caps insulated with cotton balls. The number of nymphs produced was counted after 7 days.

### Making AAF1 Antibody, Co-IP, and Immunoblot Analysis

A peptide antibody against AAF1 (peptide sequence: DRQTVRRVKTVKRVNQV) was produced and purified by Alpha Diagnostic International Inc. Immunoblot analysis of the Co-IP eluates and total leaf proteins was performed as previously described ^132^. Briefly, for Co-IP experiments, 30 µl of ChromoTek GFP-Trap Magnetic Particles M-270 or agarose beads (Proteintech) was added to 2 ml of cleared leaf lysate and incubated for 4 h at 4 °C. After washing, samples were eluted with 50 µl of 2× Laemmli sample buffer, boiled for 5 min, and 15 µl was loaded onto SDS-PAGE gels. Proteins were transferred to PVDF membranes and probed with the following antibodies: anti-GFP (BioLegend, Cat: 902605 was; 1:10,000), anti-GFP (Roche, Cat: 11814460001; 1:2,000), anti-mCherry (Mouse Monoclonal 8E5.G7 IgG2a kappa, ROCKLAND; 1:5,000), anti-AAF1 (1:5,000), anti-ubiquitin (UBQ11, Agrisera; 1:10,000), anti-COXII (AS04053A, Agrisera; 1:10,000), and anti-catalase (AS09501, Agrisera; 1:5,000). The anti-GFP antibody from BioLegend (Cat. No. 902605) was used for all GFP blots unless otherwise specified; in those cases, the antibody from Roche™ (Cat. No. 11814460001) was used. If Co-IP eluates were intended for LC-MS, beads were washed and proteins were eluted with 150 µl of 25 µM HCl, followed by neutralization with 20 µl of Tris (pH 8.0) prior to LC-MS analysis.

### Lipid Analysis

Leaf punches were collected in liquid nitrogen and stored at −80 °C and on dry ice until lipid extraction at the Kansas Lipidomics Research Center Analytical Laboratory, Kansas State University. For lipid extraction, samples were immersed in 0.4 ml of pre-heated isopropanol (0.02% BHT), incubated at 60 °C for 30 minutes, and then allowed to cool. Subsequently, 1.2 ml of a chloroform/methanol/water mixture (30:41.5:3.5, v/v/v) was added to reach a final volume of 1.6 ml. Samples were mixed and shaken at 50–100 rpm for 24 hours. For LC-MS analysis, solvents were transferred to clean vials. The analysis was performed using multiple reaction monitoring with direct infusion on a triple quadrupole mass spectrometer equipped with an electrospray ionization source, as previously described ^134^, using an internal standard mixture. For this study, the method was adapted to a Sciex 6500+ mass spectrometer, and the data were processed using the LipidomeDB Data Calculation Environment ^135^. Raw data are provided in Data Sets S4 and 5. These datasets also include measurements from an additional replicate set of plants grown in soil without Osmocote fertilizer prior to lipid analysis (labeled as −P samples). Although these data were not included in the final analyses, they are provided in the raw datasets for completeness.

### Assay of relative electrolyte leakage

The relative electrolyte leakage assay was used as an indicator of cell death–derived membrane damage. Briefly, three 0.5-cm leaf discs from each leaf sample were collected using a leaf punch, washed in deionized water, and then immersed in 2 mL of deionized water while rotating at 26 °C. The rotation time was 3 h or 2 h for plants expressing GFP–AAF1 or HA–AAF1, respectively. Conductivities were measured with a pocket conductivity detector Horiba LAQUAtwin EC-11 Compact Conductivity Meter, # 3999960125 (ABQ Industrial, Texas, USA). The assay was done on leaves from six different plants as biological replicates.

### RNA Extraction, and RT-qPCR analysis

Total RNA was isolated from leaf tissues using the RNeasy Plant Mini Kit (Qiagen™, Cat. No. 74904) according to the manufacturer’s instructions. One microgram of total RNA was reverse transcribed into cDNA using the High-Capacity cDNA Reverse Transcription Kit (Applied Biosystems™, Cat. No. 4368814). The resulting cDNA was used as a template for quantitative PCR (qPCR) with gene-specific primers. RT-qPCR was performed using Applied Biosystems™ SYBR™ Green Universal Master Mix (Cat. No. 43-091-55) on an ABI PRISM™ 7900HT Sequence Detection System (Applied Biosystems, CA, USA). Relative gene expression levels were normalized against two reference genes, as previously described ^136^. Primer sequences for the reference genes *Actin* and *PP2A* (*protein phosphatase 2A*) were obtained from reference ^137^, while primers for target genes were designed in this study. All primer sequences are listed in Data Set S7. Each tissue sample included three biological replicates, with each biological replicate analyzed using three technical replicates.

### Statistical analysis

t-tests and ANOVA were performed using built-in functions in Microsoft Excel. Chi-square and Tukey’s HSD tests were calculated manually in Excel. Dunnett’s tests were performed using R. Raw data and statistical analyses for all bar graphs and line graphs are presented in Supplementary Data Set S1.

## Supporting information

Feiz et al Supplemental Figures

## Acknowledgements

The authors would like to thank Dr. Céline Masclaux-Daubresse for providing the *atg5* Ubi:GFP-ATG8a seeds, Dr. Corinne Schmitt-Keichinger for providing the set of modified pEAQ vectors, Dr. Maria Harrison for providing the pK7 m34GW destination vector and constructs for expression of GFP and mCherry–HDEL, and Dr. Ruth Welti and Mary Roth for performing the LC-MS analysis of the lipids.

## Author Contributions

L.F. designed research, performed experiments, analyzed data, oversaw the project, and wrote the paper. N.M. performed research and analyzed data (LC–MS analyses of the aphid salivary proteome and Co-IP samples). H.F. performed research and analyzed data (VIGS-based aphid bioassays). S.I. performed research (cloning genes into the pCAMBIA2300 double-expression vector). V.T. and D.A. performed research and analyzed data (LC–MS proteomic analyses of Arabidopsis and *N. benthamiana* leaf tissues). G.J. performed research and secured funding. All authors read the manuscript and provided feedback.

---

**Video 1a**. (In support of Fig. S15a). Movie captured by sequential confocal scanning showing dynamic spherical structures in *N. benthamiana* leaf cells marked by complemented interaction between nYFP–AAF1 and cYFP–RABD2a (green fluorescence). GUS–mCherry (red fluorescence) was co-expressed from the same T-DNA as a cytosolic transformation marker. The movie is an overlay of green, red, and bright-field channels. Scale bar = 100 μm.

**Video 1b**. (In support of Fig. S15a). Close-up movie of a different leaf region than Video 1a, captured by sequential confocal scanning, showing dynamic spherical structures in *N. benthamiana* leaf cells marked by complemented interaction between nYFP–AAF1 and cYFP–RABD2a (green fluorescence). GUS–mCherry (red fluorescence) was co-expressed from the same T-DNA as a cytosolic transformation marker. The movie is an overlay of green, red, and bright-field channels. Scale bar = 50 μm.

**Video 2**. (In support of Fig. 5a). Movie captured by sequential confocal scanning showing mobile compartments marked by the complemented interaction between nYFP–AAF1 and cYFP–RABD2a (green fluorescence). Nuclei (red fluorescence) were labeled with RFP–GTE4, a nuclear marker expressed in *N. benthamiana* cells, to indicate the relative size and position of AAF1–RABD2a–containing compartments. The movie is an overlay of green, red, and bright-field channels. Cytosolic GUS–mCherry (red fluorescence) is present at very low levels and is not detectable above background. Scale bar = 100 µm.

**Video 3a**. (In support of Fig. 5b). Merged movie of concurrent time-lapse sequences captured by sequential confocal scanning, showing mobile endosome-like organelles marked by the complemented interaction between nYFP–AAF1 and cYFP–RABD2a (green fluorescence) and autophagosomes labeled with mCherry–ATG8d (red fluorescence), all expressed in *N. benthamiana* cells. The movie is an overlay of the YFP and mCherry channels (yellow where signals overlap) and the bright-field channel. Cytosolic GUS–mCherry (red fluorescence) is present at very low levels and is not detectable above background. Scale bar = 100 µm.

**Video 3b**. (In support of Fig. 5b). Movie showing only the mCherry channel from the concurrent time-lapse sequences captured by sequential confocal scanning (merged in Video 3a). The movie shows autophagosomes labeled with mCherry–ATG8d (bright red fluorescence) associated with and trafficking on mobile endosome-like organelles (faint red fluorescence). Scale bar = 100 µm.

**Video 3c**. (In support of Fig. 5b). Movie showing only the YFP channel from the concurrent time-lapse sequences captured by sequential confocal scanning (merged in Video 3a). The movie shows mobile endosome-like organelles marked by the complemented interaction between nYFP–AAF1 and cYFP–RABD2a (green fluorescence). Scale bar = 100 µm.

**Video 4a**. (In support of Figs. S15b and 5b). Close-up merged movie of concurrent time-lapse sequences captured by sequential confocal scanning from a different leaf region than that shown in Videos 3a–c, showing a mobile endosome-like organelle marked by the complemented interaction between nYFP–AAF1 and cYFP–RABD2a (green fluorescence) and autophagosomes labeled with mCherry–ATG8d (red fluorescence), all expressed in *N. benthamiana* cells. The movie is an overlay of the YFP and mCherry channels (yellow where the signals overlap) and the bright-field channel. Cytosolic GUS–mCherry (red fluorescence) is present at very low levels and is not detectable above background. Scale bar = 50 µm.

**Video 4b**. (In support of Figs. S15b and 5b). Movie showing only the mCherry channel from the concurrent time-lapse sequences captured by sequential confocal scanning (merged in Video 4a). The movie shows autophagosomes labeled with mCherry– ATG8d (bright red fluorescence) associated with and trafficking on a mobile endosome-like organelle (faint red fluorescence). Scale bar = 50 µm.

**Video 4c**. (In support of Figs. S15b and 5b). Movie showing only the YFP channel from the concurrent time-lapse sequences captured by sequential confocal scanning (merged in Video 4a). The movie shows a mobile endosome-like organelle marked by the complemented interaction between nYFP–AAF1 and cYFP–RABD2a (green fluorescence). Scale bar = 50 µm.

**Video 5a**. (In support of Fig. 5b). Merged movie of concurrent time-lapse sequences captured by sequential confocal scanning, showing mobile endosome-like organelles marked by the complemented interaction between nYFP–AAF1 and cYFP–RABD2a (green fluorescence) and the ER labeled with mCherry–HDEL (red fluorescence), all expressed in *N. benthamiana cells*. The movie is an overlay of the YFP and mCherry channels (yellow where the signals overlap) and the bright-field channel. Cytosolic GUS–mCherry (red fluorescence) is present at very low levels and is not detectable above background. Scale bar = 75 µm.

**Video 5b**. (In support of Fig. 5b). Movie showing only the mCherry channel from the concurrent time-lapse sequences captured by sequential confocal scanning (merged in Video 5a). The movie shows the ER labeled with mCherry–HDEL exchanging material with mobile endosome-like organelles (dynamic red fluorescence). Scale bar = 75 µm.

**Video 5c**. (In support of Fig. 5b). Movie showing only the YFP channel from the concurrent time-lapse sequences captured by sequential confocal scanning (merged in Video 5a). The movie shows mobile endosome-like organelles marked by the complemented interaction between nYFP–AAF1 and cYFP–RABD2a (green fluorescence). Scale bar = 75 µm.

**Video 6a**. (In support of Fig. 5b). Merged movie of concurrent time-lapse sequences captured by sequential confocal scanning from a different leaf region than that shown in Videos 5a–c, showing mobile endosome-like organelles marked by the complemented interaction between nYFP–AAF1 and cYFP–RABD2a (green fluorescence) and the ER labeled with mCherry–HDEL (red fluorescence), all expressed in *N. benthamiana cells*. The movie is an overlay of the YFP and mCherry channels (yellow where the signals overlap). Cytosolic GUS–mCherry (red fluorescence) is present at very low levels and is not detectable above background. Scale bar = 75 µm.

**Video 6b**. (In support of Fig. 5b). Movie showing only the mCherry channel from the concurrent time-lapse sequences captured by sequential confocal scanning (merged in Video 6a). The movie shows the ER labeled with mCherry–HDEL exchanging material with mobile endosome-like organelles (dynamic red fluorescence). Scale bar = 75 µm.

**Video 6c**. (In support of Fig. 5b). Movie showing only the YFP channel from the concurrent time-lapse sequences captured by sequential confocal scanning (merged in Video 6a). The movie shows mobile endosome-like organelles marked by the complemented interaction between nYFP–AAF1 and cYFP–RABD2a (green fluorescence). Scale bar = 75 µm.

## Supplementary Data and Figure Legends

**Data Set 1. (**In support of Figs. 1, 3, 4, 6, S4, S9, S14, S16, and S19). (Raw data analysis).

**Data Set 2. (**In support of Figs. 2, 4, and 5). Sorted list of identified proteins and their corresponding peptides from the Co-IP of GFP-AAF1.

**Data Set 3.** (In support of Fig. 3g and Table S1). Quantitative proteomics data from *N. benthamiana* tissues.

**Data Set 4.** (In support of Figs. 4 and S14). Lipidomic data.

**Data Set 5. (**In support of Figs. 4 and S14). Lipidomic data.

**Data Set 6.** (In support of Figs. 7b and S17 and Table S2). Quantitative proteomics data from Arabidopsis tissues.

**Data Set 7. (**In support of all Figs.). Primers and constructs.

**Fig. S1.** (In support of Fig. 1). *Agrobacterium tumefaciens-*mediated transient expression of 48 secreted aphid salivary peptides in *N. benthamiana* leaves. (**a**) Schematic representation of gene cassettes encoding small salivary proteins (10–25 kDa) after removal of their signal peptides, which were cloned into expression vectors and introduced into *A. tumefaciens*. (**b**) Transient expression in *N. benthamiana* leaf cells. Arrows indicate leaves expressing AAF1 (SP11 in this figure), along with SP15 and SP37, which served as negative controls.

**Fig. S2. (**In support of Figs. 1, 2, 3, 5 and 6). The expression of AAF1 fused to different tags at varying levels from different vectors. (**a**) *N. benthamiana* leaf phenotypes imaged ten days after infiltration with constructs expressing GFP-AAF1, AAF1-HA(3x), and HA-AAF1. (**b**) Immunoblot analysis of total proteins extracted from the leaves shown in (**a**). Blots were probed with an AAF1-specific antibody; arrows indicate bands corresponding to AAF1 fused to different tags. The top and middle panels show the same blot with different exposures. Ponceau staining was used as a loading control.

**Fig. S3.** (In support of Figs. 1, 2, 3, and 5). Alignment of AAF1 from different aphid species. T-Coffee (https://tcoffee.crg.eu/apps/tcoffee/do:regular) was used to generate a FASTA-aligned file from the aphid sequences, followed by alignment shading using Boxshade (https://junli.netlify.app/apps/boxshade/#forms::boxshade). The green rectangle indicates the secretion signal peptide, the red rectangle marks the position of a putative AIM motif, and the blue rectangle shows the location of the Rab-binding domain. The GenBank accession number associated with the protein sequence from each aphid species is shown in the bottom right corner.

**Fig. S4.** (In support of Fig. 1). Catalase1 overexpression reduces AAF1-mediated cell death, H₂O₂ production, and aphid reproduction. (**a**) H₂O₂ levels measured in *N. benthamiana* leaves expressing GFP, RFP-AAF1, or RFP-AAF1 + Catalase-GFP. Data represent the mean ± s.e. (*N* = 3). Statistical analysis was performed using one-way ANOVA followed by Tukey’s HSD test; different letters indicate significant differences (*P* < 0.05). (**b**) *N. benthamiana* leaf phenotypes 10 days post-infiltration with constructs expressing RFP-AAF1, RFP-AAF1 + Catalase-GFP, or Catalase-GFP alone. (**c**) Aphid reproduction on *asat2 N. benthamiana* leaves transiently expressing Catalase-GFP or GFP (control), using one original nymph per cage, one cage per plant. Data represent the mean ± s.e. (*N* = 20); \*\**P* < 0.01, *t*-test. (d) Aphid survivorship on *asat2 N. benthamiana* leaves transiently expressing Catalase-GFP or GFP (control), using ten original nymph per cage, two cages per plant. Data represent the mean ± s.e. (*N* = 16 for GFP, and 17 for Catalase-GFP); \**P* < 0.05, *t*-test.

**Fig. S5.** (In support of Figs. 1d-e). GFP-AAF1–induced symptoms are alleviated in *N. benthamiana* leaves treated with VIGS-mediated RNAi targeting *AAF1*, but not in leaves treated with the TRV2 empty vector, confirming effective silencing of AAF1 in the former. Images were taken one week after GFP-AAF1 expression and three weeks after infiltration with either Vi-*AAF1* (expressing *AAF1* RNAi) or the Vi-TRV2 empty vector control.

**Fig. S6.** (In support of Figs. 2 and 5). RABD1 and RABD2a interact with AAF1. Independent replicates of the Co-IP assays shown in Fig. 2a were performed using AAF1 fused to different C-terminal tags—AAF1-HA(3x) (**a**) or AAF1-FLAG (**b**)—co-expressed with the same GFP-tagged bait proteins used in Fig. 2a (free GFP, GFP-RABD1, GFP-RABD2a, or GFP-ATG8f) in *N. benthamiana* leaves. A GFP-Trap Co-IP assay was conducted to pull down AAF1 using the GFP-fused baits. Immunoblots of the Co-IP eluates and inputs were probed with anti-AAF1 and anti-GFP antibodies. (**c**) Subcellular co-localization of GFP-AAF1 with RFP-RABD1 in leaves was analyzed using confocal microscopy, with GFP–AAF1 shown in green and RFP–RABD1 in red; their overlap indicates colocalization in yellow. White arrowheads indicate punctate structures; scale bar = 10 µm.

**Fig. S7.** (In support of Figs. 2, 3, and 5). Structural modeling of AAF1 predicted a C-terminal Rab-binding domain. (a) Phyre2 was used to model the structure of six AAF1 protein segments. Three segments, including segment 6, which is homologous to the Rab-binding domain of a guanine nucleotide exchange factor (GEF) that acts as an activator and binding partner of the yeast RAB1 (Ypt1), showed moderate to high confidence scores in structural homology. (b) Structural modeling of segment 6 using I-TASSER confirmed its homology to the Rab-binding domain of the same yeast protein, as well as to the Rab-binding domain of a human RAB-activating protein.

**Fig. S8.** (In support of Fig. 2). Subcellular localization of (a) GFP-AAF1 or (b) GFP-Frag-1, co-expressed with the ER marker mCherry-HDEL in *N. benthamiana* leaves, was analyzed using confocal microscopy. Two individual frames from the z-stack, as well as the full z-stack, are shown. Insets show 4.5× magnified views of the indicated areas. White arrows indicate punctate structures, and open triangle indicate the endoplasmic reticulum surrounding the nucleus. Scale bar = 10 µm. (c) A replicate of the Co-IP eluate blot shown in Fig. 2h, which also included a GFP-only negative control as bait, was probed with an anti-ubiquitin antibody.

**Fig. S9.** (In support of Figs. 2-4). AAF1 colocalizes with ATG8d in autophagosomes. Transient co-expression of GFP-AAF1 with mCherry-ATG8d (**a**, top panel) and RFP-AAF1 with GFP-ATG8d (a, bottom panel) was performed in *N. benthamiana* leaves. Confocal microscopy reveals single optical segments, with GFP-AAF1 and GFP-ATG8d shown in green, mCherry-ATG8d and RFP-AAF1 in red, and their overlap indicating colocalization in yellow. White arrowheads mark punctate structures where GFP and RFP signals coincide. Scale bar = 10 µm. (**b**, **c**) Intensity plots show relative GFP, RFP, or mCherry fluorescence signals along randomly drawn lines spanning GFP–AAF1-, RFP–RABD1 (from Fig. S7c)-, or GFP–AAF1-, mCherry–ATG8d (**a**)-marked puncta. Representative plots demonstrate that GFP–AAF1 fluorescence peaks overlap more strongly with RFP–RABD1 (Fig. S6c) than with mCherry–ATG8d (**a**). (**d**) Bar charts show the percentage of GFP–AAF1 puncta that colocalize with either RFP–RABD1 or mCherry–ATG8d. Data represent 47 individual puncta quantified from at least 10 images obtained from independent biological replicates (raw data provided in Dataset S1). A chi-square test revealed a significant difference in the frequency of GFP–AAF1 colocalization with RFP–RABD1 versus mCherry–ATG8d (\**P* < 0.01). (**e**) Z-stack and two single-plane images from an independent experiment coexpressing GFP–AAF1 and mCherry–ATG8d.

**Fig. S10.** (In support of Fig. 3). (**a**) AAF1 increases autophagic flux. Proteins extracted from *N. benthamiana* leaves transiently co-expressing GFP–ATG8d with or without HA– AAF1 were analyzed by immunoblotting using antibodies against GFP and AAF1. For the light treatment, plants were infiltrated 48 h prior to sampling and exposed to light for 6 h on day 3 post-infiltration. In the dark treatment, plants were kept in darkness during this period following the standard overnight dark cycle. (**b**) As in (**a**), *N. benthamiana* leaves transiently co-expressing HA–AAF1 or HA–Frag2 with GFP–ATG8d were analyzed by immunoblotting using antibodies against GFP and AAF1. (**c**) Aphids were caged for 24 and 48 h on *N. benthamiana* leaves transiently expressing GFP–ATG8f at 48 h post infiltration. Leaf sections from the caged areas were analyzed by immunoblotting using antibodies against GFP. (**d**) Transient expression of AAF1 increases the number and abundance of ubiquitinated proteins. Immunoblot analysis was performed on total proteins extracted from *N. benthamiana* leaves two days after transient expression of GFP or GFP-AAF1. Equal surface areas of the leaves or their dilutions (as indicated) were analyzed using antibodies specific to the proteins listed on the right. CBB, Coomassie Brilliant Blue-stained gel, was used as the loading control. The anti-GFP antibody from BioLegend (Cat. No. 902605) was used for all GFP blots in this image.

**Fig. S11.** (In support of Fig. 3). (**a**) HA–AAF1-induced symptoms in *N. benthamiana* leaves subjected to VIGS targeting *ATG7*, *RABD1*, or *RABD2a* were compared among the VIGS lines and with HA–Frag3-induced symptoms as a negative control. Images were taken 5 days after HA–AAF1 expression and 20 days after infiltration with Vi-*rabD1*, Vi-*rabd2a*, Vi-*atg7*, or the Vi-TRV2 empty vector control. (**b**) GFP–AAF1 protein levels from the experiment presented in Fig. 3 were monitored using immunoblotting, and (**c**) confocal imaging. (**d**) Cellular ion leakage (conductivity) of leaves from VIGS plants, used as a proxy for cell death, was measured 3 days post-infiltration. Data represent means ± s.e. from six biological replicates. Statistical analyses were performed using ANOVA; different letters indicate significant differences (*P* < 0.05, ANOVA followed by Tukey’s HSD test). Raw data are provided in Data Set 1.

**Fig. S12.** (In support of Fig. 4). structural modeling of PI3PT predicted a lipid-binding/transport protein. (**a**) Molecular modeling of full-length PI3PT using Phyre2 identified it as a lipid transport protein, specifically a phosphatidylglycerol/phosphatidylinositol transfer protein, according to the Protein Data Bank (PDB). The structure was predicted with 100% confidence based on the crystal structure of the *S. cerevisiae* Niemann-Pick type C protein Npc2, bound to ergosterol. (**b**) Molecular modeling using I-TASSER confirmed the result obtained with Phyre2.

**Fig. S13.** (In support of Fig. 4). PI3PT interacts with AAF1. (**a**-**b**) Independent replicates of the GFP-Trap Co-IP shown in Fig. 4b were performed using GFP-tagged bait proteins: GFP and GFP-ATG8f (both negative controls), GFP-PI3PT, and GFP-RABD2a (positive control), to pull down HA-AAF1 from lysates of *N. benthamiana* leaves co-expressing each bait with HA-AAF1. Immunoblots were probed with anti-AAF1 and anti-GFP antibodies. (**c**) Overexpression of GFP-PI3PT in *N. benthamiana* leaves did not induce cell death responses similar to those caused by AAF1, RABD1, and RABD2a. Leaves were imaged 10 days post-infiltration.

**Fig. S14.** (In support of Fig. 4). AAF1 promotes the accumulation of specific lipids, including lipid droplet constituents. AAF1 expression increases the levels of (**a**) stigmasterol glucoside (Glc) esters, and (**b**) triglycerides/triacylglycerols (TGs) in plants. Total lipids were measured in *N. benthamiana* leaves transiently expressing GFP-AAF1, HA-AAF1, or GFP as a negative control. For each sample, lipid content (nmol of lipid per mg dry tissue) was measured by LC-MS, as described in the methods. Statistical analysis was performed using ANOVA; different letters indicate significant differences (*P* < 0.05, ANOVA followed by Tukey’s HSD test). Raw data are provided in Data Sets 3 and 4. These datasets also include measurements from an additional set of replicate plants, which were grown in soil without Osmocote fertilizer prior to total lipid analysis (labeled as –P samples). Although these data were not included in the final analyses, they are provided in the raw datasets for completeness. (**c**) Lipid droplets (LDs) colocalize with and accumulate near AAF1-containing punctate structures. *N. benthamiana* leaves expressing RFP-AAF1 (red puncta) two days post-infiltration were treated with 10 µM BODIPY and analyzed using confocal microscopy; scale bar = 25 µm.

**Fig. S15.** (In support of Fig. 5). (**a**) Interacting AAF1 and RABD1 or RABD2a form previously uncharacterized mobile organelles (Videos 1a-b, and 4a-c support this figure). Sequential confocal images show YFP complementation (green fluorescence) in 2–8 µm spherical structures in *N. benthamiana* leaf cells coexpressing nYFP–AAF1 and cYFP–RABD2a or cYFP–RABD1, but not in cells coexpressing nYFP–Frag3 (a truncated AAF1 lacking the Rab-binding domain; negative control) and cYFP–RABD2a. GUS–mCherry (red fluorescence) was co-expressed from the same T-DNA as a cytosolic transformation marker. Scale bars = 50 μm. (**b**) Images show an independent replicate of the first-row images in Fig. 5b, taken from a different leaf area, and depict endosome-like organelles marked by the complemented interaction of nYFP–AAF1 and cYFP–RABD2a (green fluorescence), autophagosomes marked by mCherry–ATG8d (red fluorescence), and their overlap (yellow), following co-expression from independent plasmids. Insets show 4×-magnified views of the indicated areas. Scale bars = 10 μm. In all images, the overlay panels include bright-field images.

**Fig. S16.** (In support of Fig. 6). AAF1 enhances plant growth. An independent replicate of the experiments presented in Fig. 6c, e, and h, including only three of the four genotypes (Col-0, Col-0+AAF1, and *atg5*+AAF1). (**a**) Rosette surface areas of mature (35-day-old) plants were measured using Fiji after capturing images at the same scale for each plant. Data represent the mean ± s.e. (N = 17 for *atg5*+AAF1; N = 18 for the other two genotypes). (**b**) Dry weight of the same plants. Statistical analysis was performed using ANOVA; different letters indicate significant differences (*P* < 0.001; ANOVA followed by Tukey’s HSD test). (**c**) Immunoblot analysis of total proteins extracted from leaves of the three Arabidopsis genotypes described in (**a**). Blots were probed with anti-AAF1 and anti-GFP (BioLegend) antibodies.

**Fig. S17.** (In support of Fig. 7b) Comparative LC-MS proteomic analysis of pooled seedlings harvested from the same plant batches shown in Fig. 6a and used for the immunoblot analysis in Fig. 7a. Volcano plot showing differential protein abundance between *atg5*+AAF1 and *atg5* (three biological replicates per genotype). P_only, *P*-value cutoff; ns, not significant. Differentially abundant proteins are listed in Table S2, and the complete proteomic dataset is provided in Data Set S6.

**Fig. S18.** (In support of Figs. 6 and 7). Ubi:GFP-ATG8a expression (green fluorescence) marks the presence and abundance of autophagosomes and autophagic bodies in roots of Arabidopsis lines Col-0, Col-0+AAF1, and *atg5*+AAF1. Roots from 5-day-old seedlings grown on ½ MS medium without sucrose for 5 days were treated with 1 µM concanamycin A, with or without 100 µM BTH, for three hours and subsequently analyzed by confocal microscopy. Insets show 3× magnified views of the indicated areas. Scale bars = 10 µm.

**Fig. S19.** (In support of Fig. 7e). An independent replicate of the experiment presented in Fig. 7d. Single one-day-old *M. persicae* aphid nymphs were caged on a single rosette-stage leaf of each Arabidopsis plant representing the four genotypes (Col-0, Col-0+AAF1, *atg5*+AAF1, and *atg5*), and the number of progeny produced was counted seven days later. Data represent means ± s.e. (N = 11 for Col-0+AAF1 and N = 8 for each of the other three genotypes). Statistical analysis was performed using one-way ANOVA followed by Tukey’s HSD test (*P* < 0.05).

**Fig. S20.** (In support of Fig. 1). Schematic of the proteomic analysis of aphid saliva that led to the identification of AAF1.

**Table. S1.** (In support of Fig. 3g). List of proteins in each of the seven clusters showing differential abundance among Vi-TRV2 plants expressing GFP–AAF1 (severe cell death), Vi-*rabd2a* plants expressing GFP–AAF1 (low cell death), and Vi-TRV2 plants expressing GFP–Frag2 (control; no cell death), as shown in Fig. 3g. The complete proteomic dataset is provided in Data Set S3.

**Table. S2.** (In support of Figs. 7b and S17). Proteins that were significantly up-or downregulated in pairwise comparisons between Col-0+AAF1 and Col-0 (control) and between *atg5*+AAF1 and *atg5* are listed. The complete proteomic dataset is provided in Data Set S6.

