## Supplementary material for "An Aphid Protein Interferes with Plant endomembrane Trafficking Machinery to Promote Aphid Survival and Plant Growth": Feiz et al Supplemental Figures

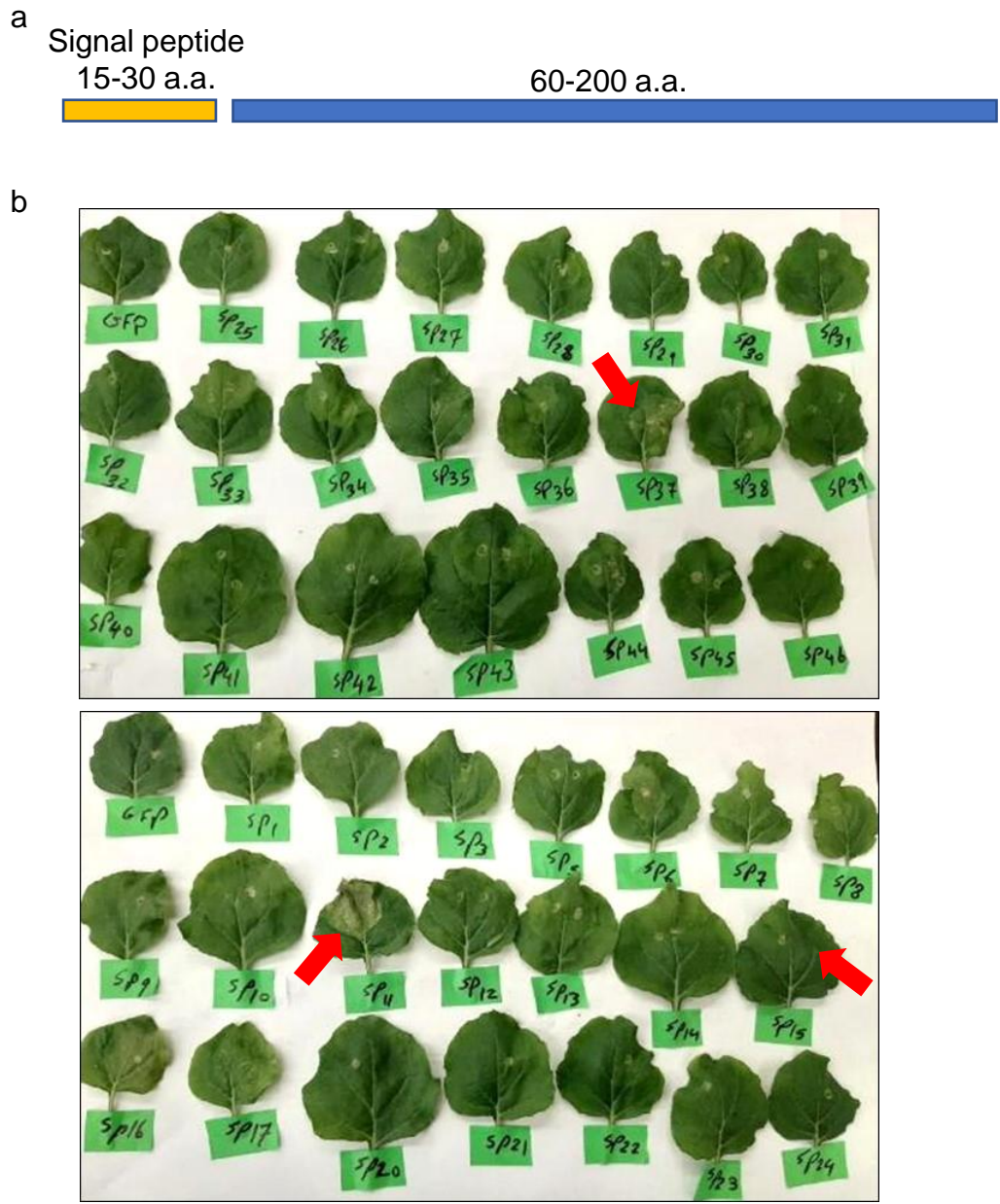

**Figure S1.** In support of Fig. 1. *Agrobacterium tumefaciens*-mediated transient expression of 48 secreted aphid salivary peptides in *Nicotiana benthamiana* leaves. (a) Schematic representation of gene cassettes encoding small salivary proteins (10–25 kDa) after removal of their signal peptides, which were cloned into expression vectors and introduced into *A. tumefaciens*. (b) Transient expression in *N. benthamiana* leaf cells. Arrows indicate leaves expressing AAF1 (SP11 in this figure), along with SP15 and SP37, which served as negative controls.

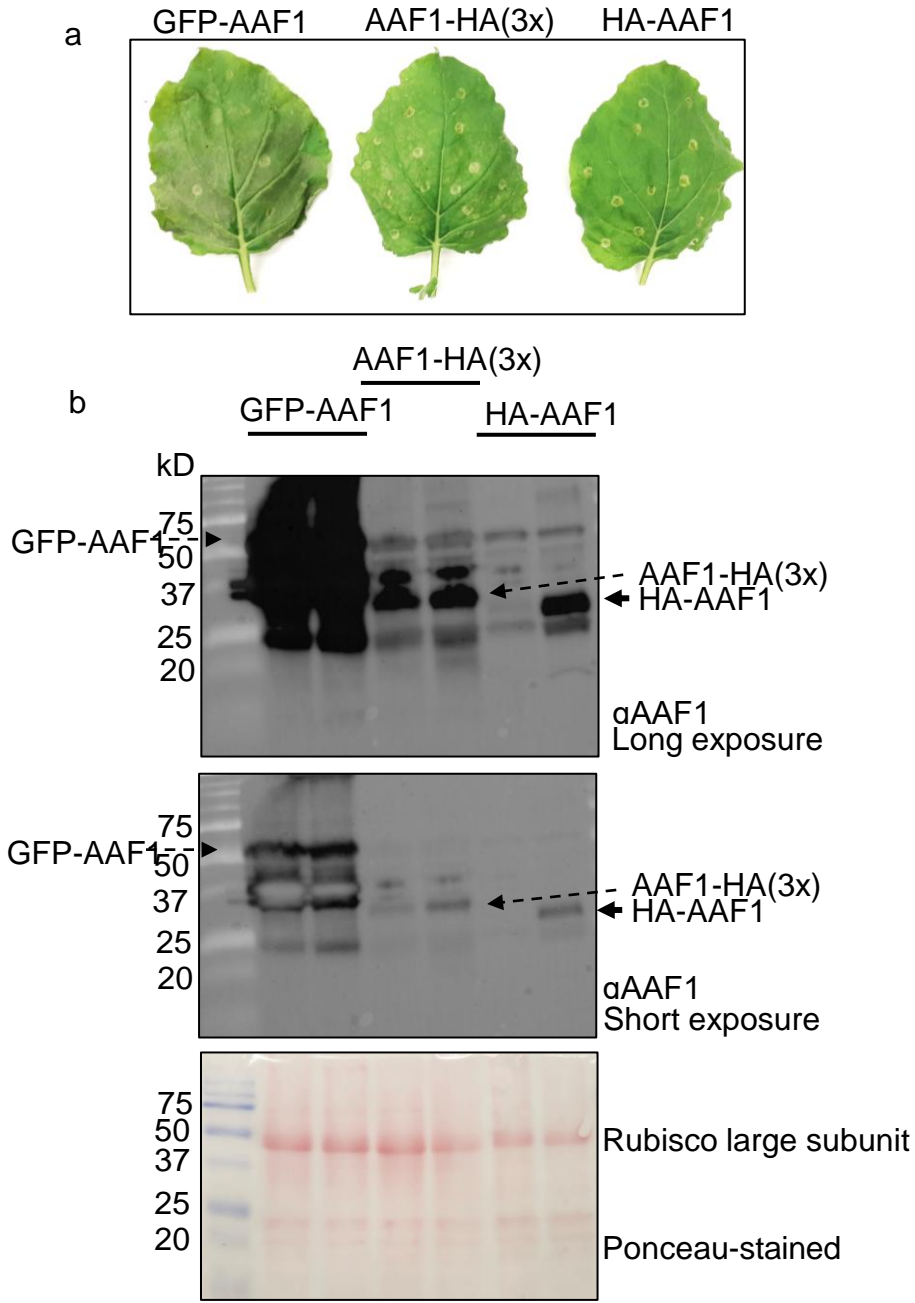

**Figure S2.** In support of Figs. 1, 2, 3, 5 and 6. The expression of AAF1 fused to different tags at varying levels from different vectors. **(a)** *N. benthamiana* leaf phenotypes imaged ten days after infiltration with constructs expressing GFP-AAF1, AAF1-HA(3x), and HA-AAF1. **(b)** Immunoblot analysis of total proteins extracted from the leaves shown in **(a)**. Blots were probed with an AAF1-specific antibody; arrows indicate bands corresponding to AAF1 fused to different tags. The top and middle panels show the same blot with different exposures. Ponceau staining was used as a loading control.

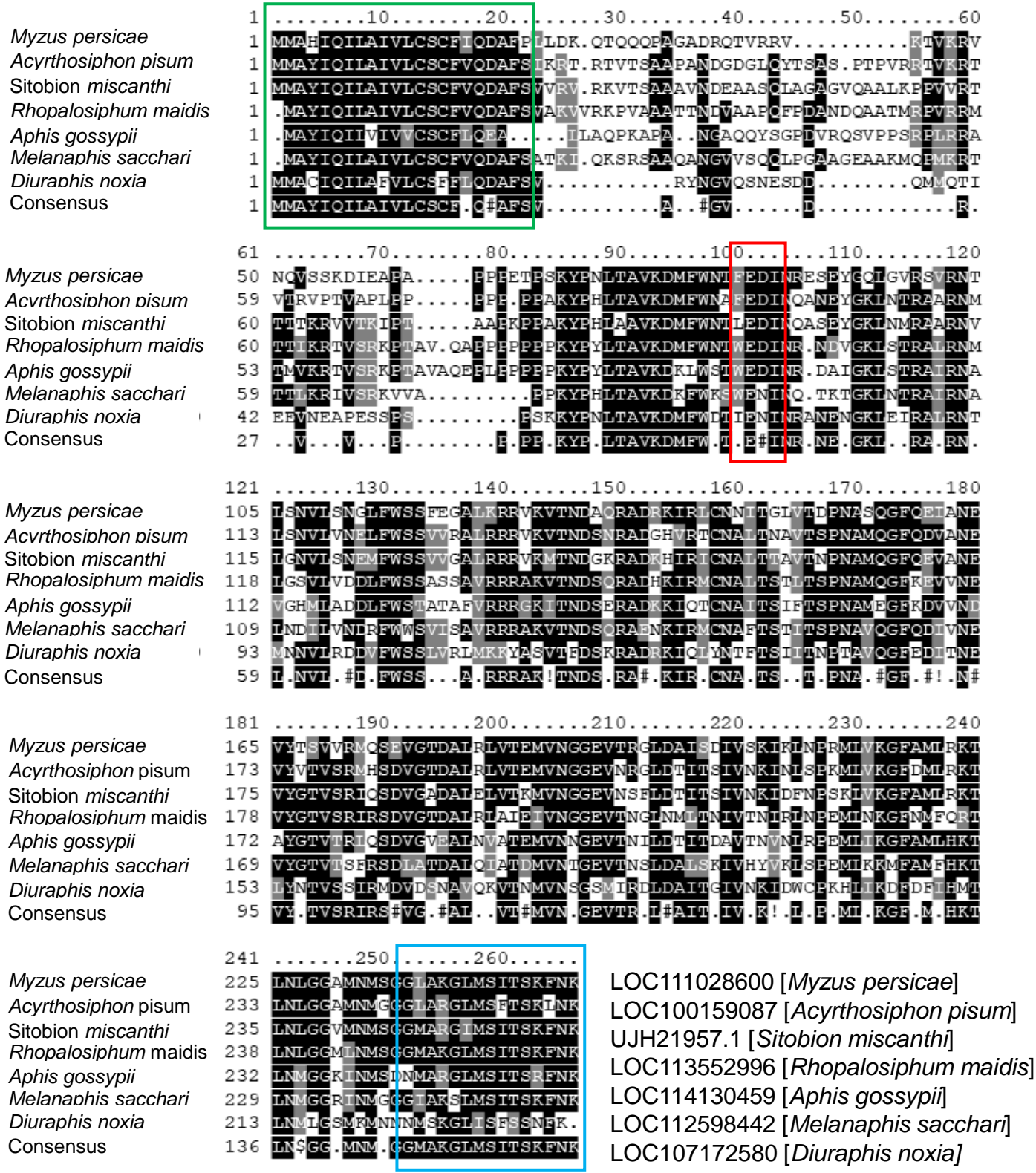

**Figure S3.** In support of Figs. 1, 2, 3, and 5. Alignment of AAF1 from different aphid species. T-Coffee (https://tcoffee.org.eu/apps/tcoffee/do:regular) was used to generate a FASTA-aligned file from the aphid sequences, followed by alignment shading using Boxshade (https://junli.netlify.app/apps/boxshade/#forms:boxshade). The green rectangle indicates the secretion signal peptide, the red rectangle marks the position of a putative AIM motif, and the blue rectangle shows the location of the Rab-binding domain. The GenBank accession number associated with the protein sequence from each aphid species is shown in the bottom right corner.

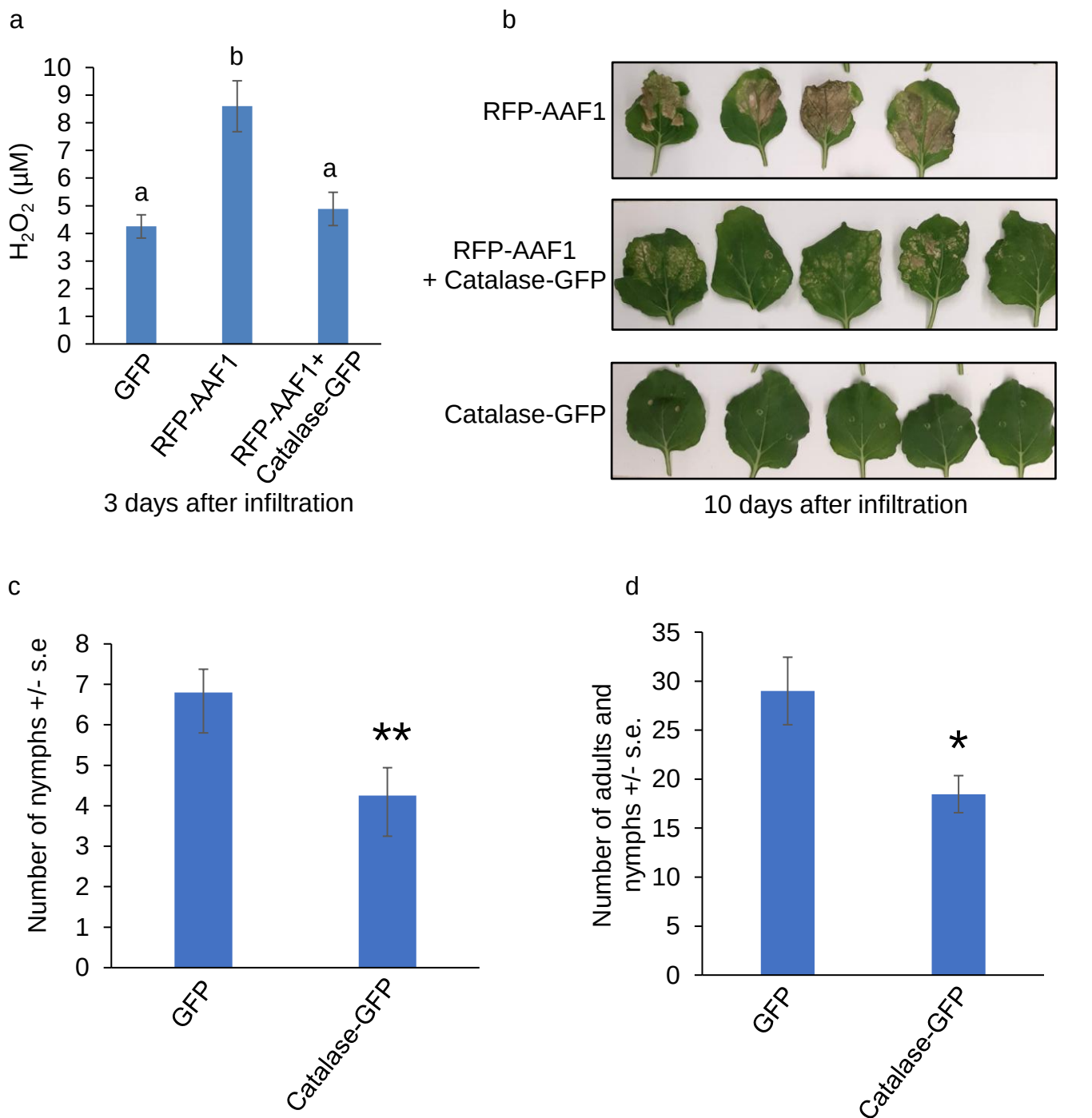

**Figure S4.** In support of Fig. 1. Catalase1 overexpression reduces AAF1-mediated cell death, H<sub>2</sub>O<sub>2</sub> production, and aphid reproduction. **(a)** H<sub>2</sub>O<sub>2</sub> levels measured in *N. benthamiana* leaves expressing GFP, RFP-AAF1, or RFP-AAF1 + Catalase-GFP. Data represent the mean ± s.e. ($N = 3$). Statistical analysis was performed using one-way ANOVA followed by Tukey's HSD test; different letters indicate significant differences ($P < 0.05$). **(b)** *N. benthamiana* leaf phenotypes 10 days post-infiltration with constructs expressing RFP-AAF1, RFP-AAF1 + Catalase-GFP, or Catalase-GFP alone. **(c)** Aphid reproduction on *asat2* *N. benthamiana* leaves transiently expressing Catalase-GFP or GFP (control), using one original nymph per cage, one cage per plant. Data represent the mean ± s.e. ($N = 20$); \*\*$P < 0.01$, $t$-test. **(d)** Aphid survivorship on *asat2* *N. benthamiana* leaves transiently expressing Catalase-GFP or GFP (control), using ten original nymph per cage, two cages per plant. Data represent the mean ± s.e. ($N = 16$ for GFP, and 17 for Catalase-GFP); \*$P < 0.05$, $t$-test.

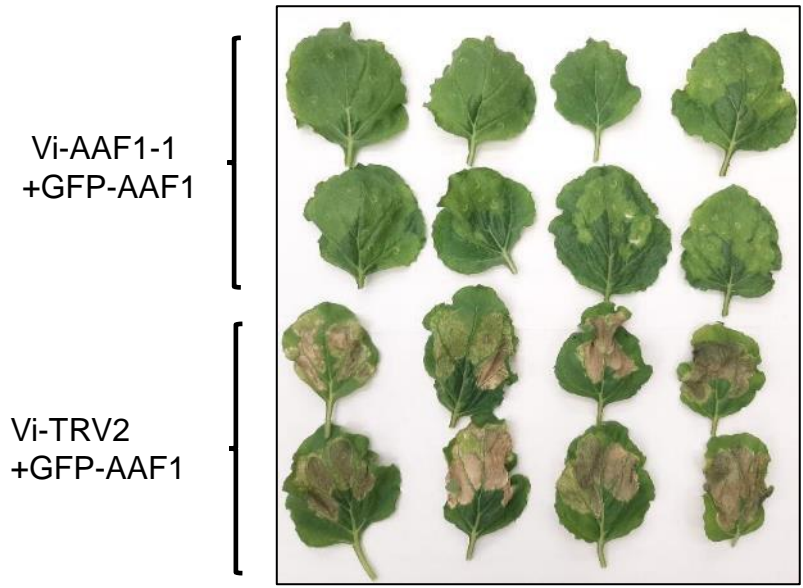

**Figure S5.** In support of Figs. 1D-E. (A) GFP-AAF1-induced symptoms are alleviated in *N. benthamiana* leaves treated with VIGS-mediated RNAi targeting *AAF1*, but not in leaves treated with the TRV2 empty vector, confirming effective silencing of *AAF1* in the former. Images were taken one week after GFP-AAF1 expression and three weeks after infiltration with either Vi-AAF1 (expressing *AAF1* RNAi) or the Vi-TRV2 empty vector control.

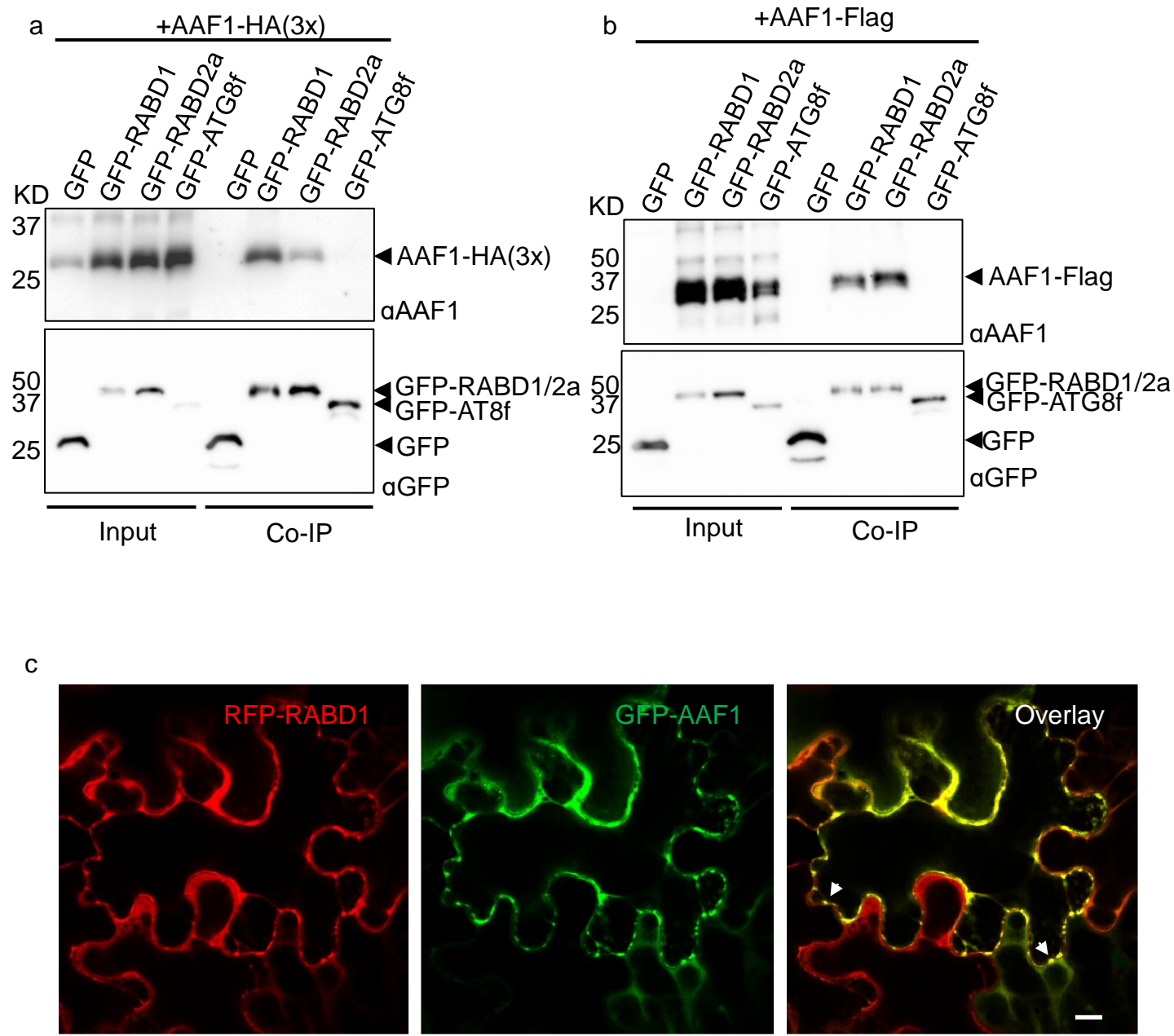

**Figure S6.** In support of Figs. 2 and 5. RABD1 and RABD2a interact with AAF1. Independent replicates of the co-immunoprecipitation (Co-IP) assays shown in Fig. 2a were performed using AAF1 fused to different C-terminal tags—AAF1-HA(3x) (**a**) or AAF1-FLAG (**b**)—co-expressed with the same GFP-tagged bait proteins used in Fig. 2a (free GFP, GFP-RABD1, GFP-RABD2a, or GFP-ATG8f) in *N. benthamiana* leaves. A GFP-Trap Co-IP assay was conducted to pull down AAF1 using the GFP-fused baits. Immunoblots of the Co-IP eluates and inputs were probed with anti-AAF1 and anti-GFP antibodies. (**c**) Subcellular co-localization of GFP-AAF1 with RFP-RABD1 in leaves was analyzed using confocal microscopy, with GFP-AAF1 shown in green and RFP-RABD1 in red; their overlap indicates colocalization in yellow. White arrowheads indicate punctate structures; scale bar = 10 μm.

a

### Segment 1=

LLDKQTQQPAGADRQTVRRVKTVKRVNQVSSKDI
EAPAPPETPSKYPN

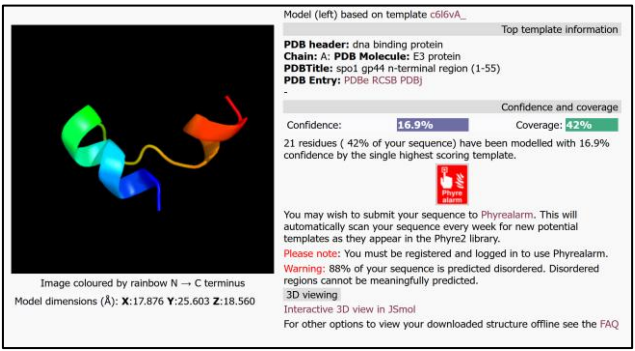

### Segment 2=

SKYPNLTAVKMFWNTFEDINRESEYGQLGVRS

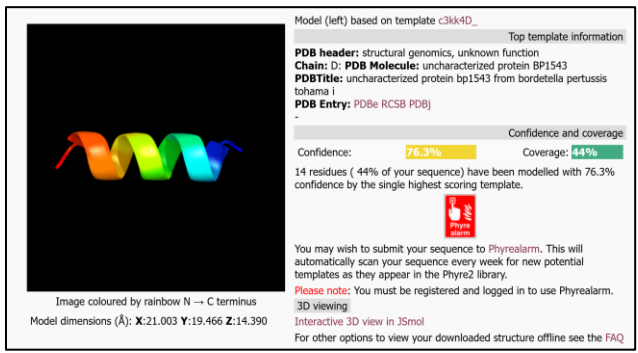

### Segment 3=

VRNTLSNVLSNGLFWSSFEGALKRRVKVTNDAQRA
DRKI

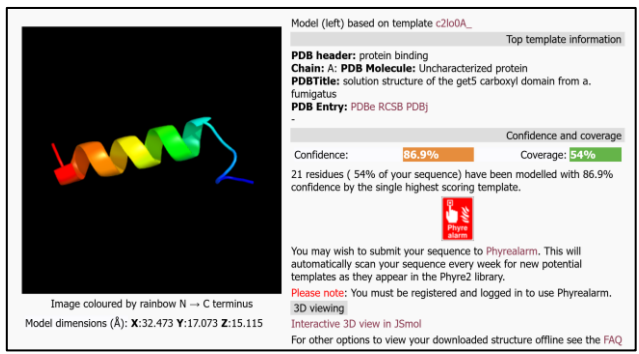

### Segment 4=

RLCNNITGLVTDPNASQGFQEIANEVYTSVVRMQ
SEVGT

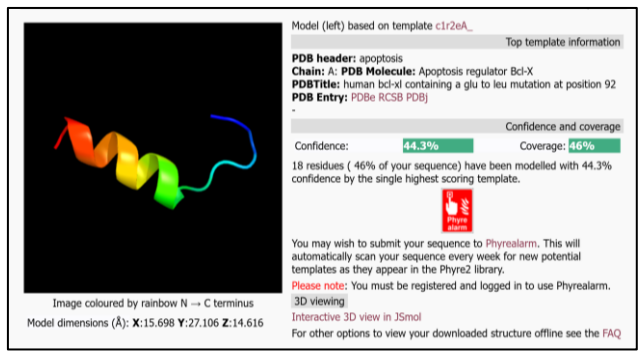

### Segment 5=

DALRLVTEMVNGGEVTRGLDAISDIVSKIKNPRMLV
KGFAM

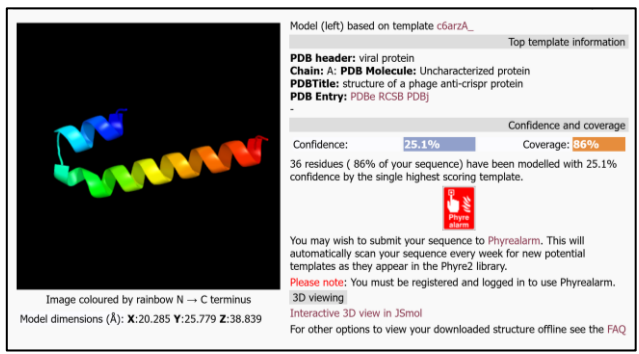

### Segment 6=

LRKTLNLGGAMNMSGGLAKGLMSITSKFNK

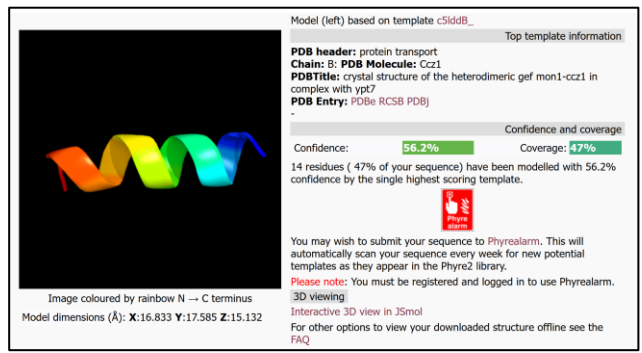

b

Segment 6=

LRKTLNLGGAMNMSGGLAKGLMSITSKFNK

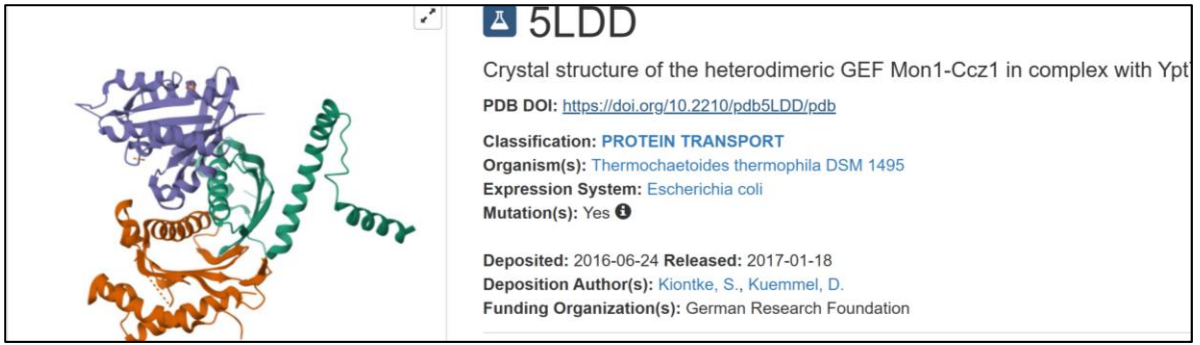

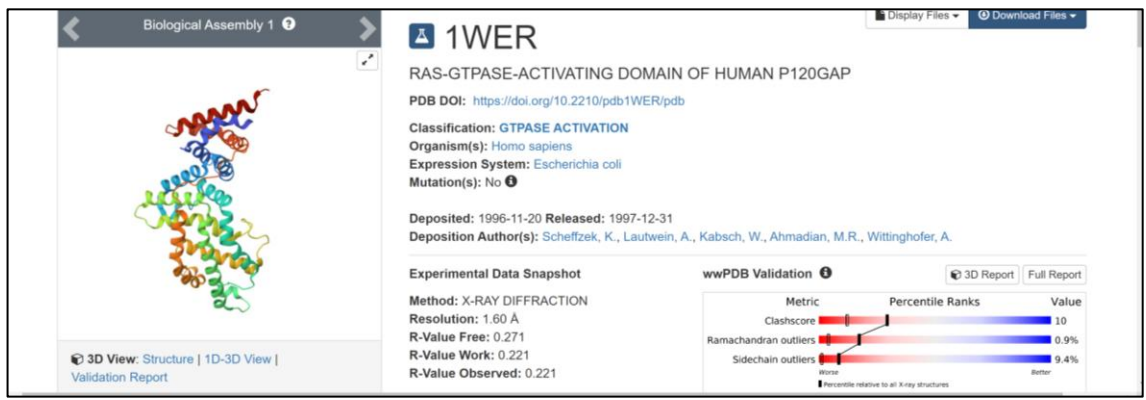

**Figure S7.** In support of Figs. 2, 3, and 5. Structural modeling of AAF1 predicted a C-terminal Rab-binding domain. (a) Phyre2 was used to model the structure of six AAF1 protein segments. Three segments, including segment 6, which is homologous to the Rab-binding domain of a guanine nucleotide exchange factor (GEF) that acts as an activator and binding partner of the yeast RAB1 (Ypt1), showed moderate to high confidence scores in structural homology. (b) Structural modeling of segment 6 using I-TASSER confirmed its homology to the Rab-binding domain of the same yeast protein, as well as to the Rab-binding domain of a human RAB-activating protein.

a

Z-9

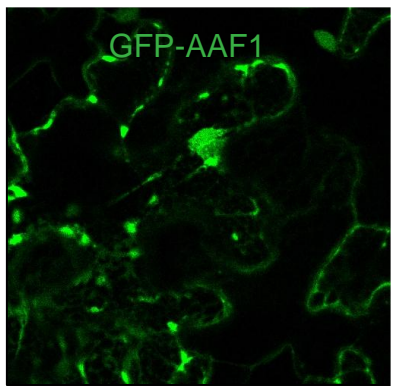

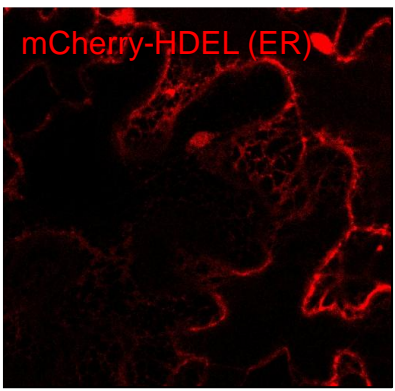

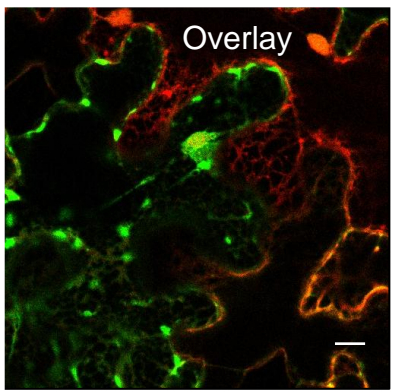

Z-24

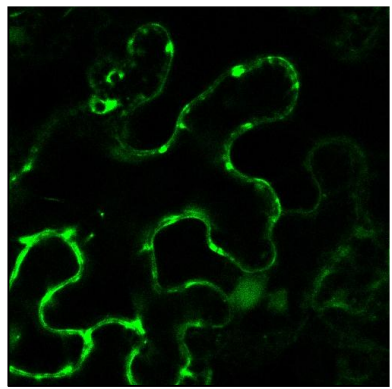

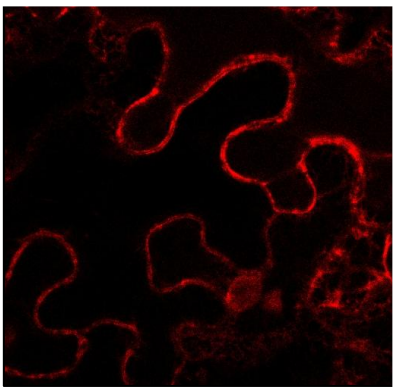

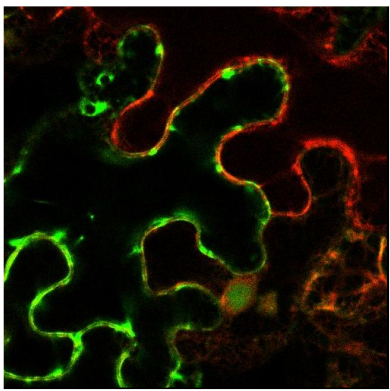

Z-stack

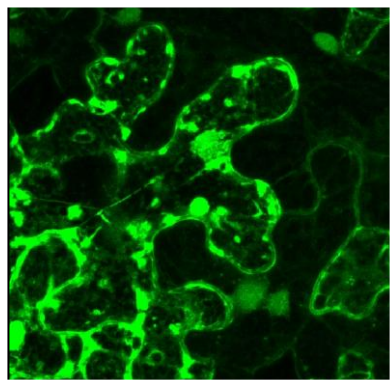

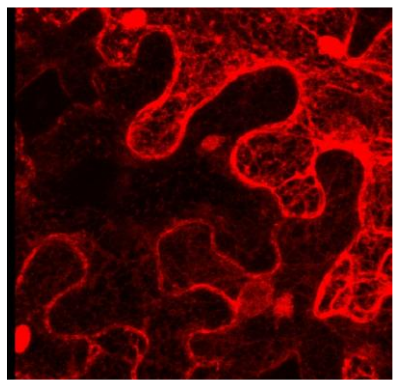

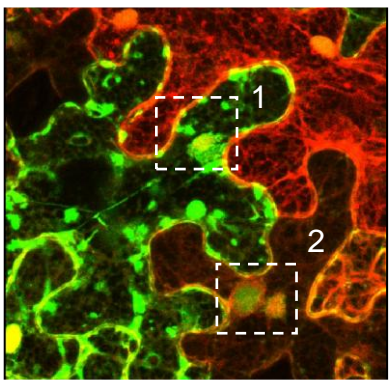

Inset 1
(4.5X)

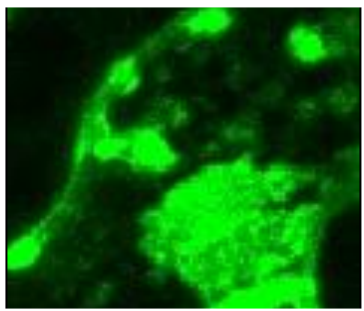

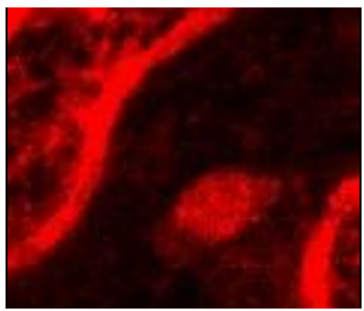

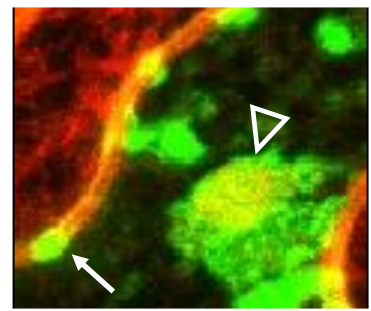

Inset 2
(4.5X)

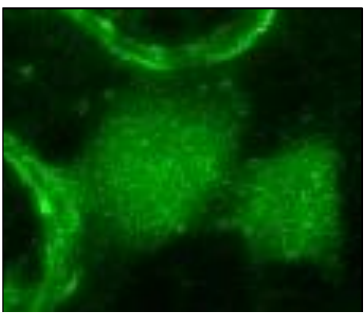

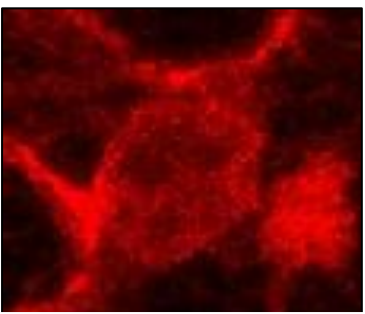

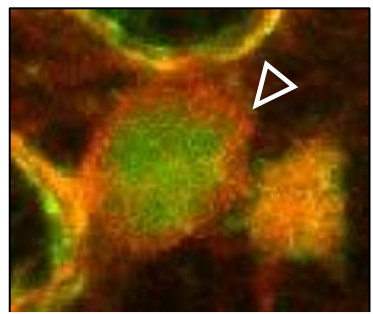

b

Z-26

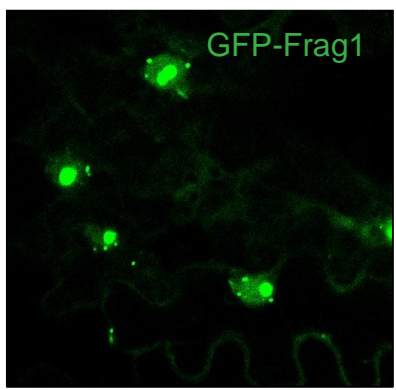

Z-33

Z-stack

Inset 1
(4.5X)

Inset 2
(4.5X)

c

**Figure S8.** In support of Fig. 2. Subcellular localization of (a) GFP-AAF1 or (b) GFP-Frag-1, coexpressed with the ER marker mCherry-HDEL in *N. benthamiana* leaves, was analyzed using confocal microscopy. Two individual frames from the z-stack, as well as the full z-stack, are shown. Insets show 4.5× magnified views of the indicated areas. White arrows indicate punctate structures, and open triangle indicate the endoplasmic reticulum surrounding the nucleus. Scale bar = 10 μm. (c) A replicate of the Co-IP eluate blot shown in Fig. 2h, which also included a GFP-only negative control as bait, was probed with an anti-ubiquitin antibody.

**Figure S18.** In support of Fig. 6. Ubi:GFP-ATG8a expression (green fluorescence) marks the presence and abundance of autophagosomes and autophagic bodies in roots of Arabidopsis lines Col-0, Col-0+AAF1, and *atg5*+AAF1. Roots from 5-day-old seedlings grown on $\frac{1}{2}$ MS medium without sucrose for 5 days were treated with 1 μM concanamycin A, without (**a**) or with 100 μM BTH (**b**), for three hours and subsequently analyzed by confocal microscopy. Insets show 3× magnified views of the indicated areas. Scale bars = 10 μm.

**Figure S19.** (In support of Fig. 7e). An independent replicate of the experiment presented in Fig. 7d. Single one-day-old *M. persicae* aphid nymphs were caged on a single rosette-stage leaf of each *Arabidopsis* plant representing the four genotypes (Col-0, Col-0+AAF1, *atg5*+AAF1, and *atg5*), and the number of progeny produced was counted seven days later. Data represent means ± s.e. (N = 11 for Col-0+AAF1 and N = 8 for each of the other three genotypes). Statistical analysis was performed using one-way ANOVA followed by Tukey's HSD test ($P < 0.05$).

*Myzus persicae*
(green peach aphid)
on *N. tabacum* leaf

Aphid diet between
two layers of Parafilm

LC-MS

Proteome

>1000 salivary
proteins,
~120 have signal
peptides → AAF1

**Figure S20.** In support of Fig. 1. Schematic of the proteomic analysis of aphid saliva that led to the identification of AAF1.
